# Development of covalent chemogenetic K_2P_ channel activators

**DOI:** 10.1101/2023.10.15.561774

**Authors:** Parker E. Deal, Haerim Lee, Abhisek Mondal, Marco Lolicato, Philipe Ribeiro Furtado de Mendonca, Holly Black, Xochina El-Hilali, Clifford Bryant, Ehud Y. Isacoff, Adam R. Renslo, Daniel L. Minor

**Affiliations:** Cardiovascular Research Institute; Department of Pharmaceutical Chemistry; Department of Molecular and Cell Biology; Helen Wills Neuroscience Institute; Weill Neurohub University of California, Berkeley, Berkeley, California 94720, United States; Molecular Biophysics and Integrated Bio-imaging Division Lawrence Berkeley National Laboratory, Berkeley, CA 94720 USA; Departments of Biochemistry and Biophysics, and Cellular and Molecular Pharmacology; California Institute for Quantitative Biomedical Research; Kavli Institute for Fundamental Neuroscience University of California, San Francisco, California 93858-2330 USA

**Keywords:** K_2P_ channel, chemogenetics, X-ray crystallography, electrophysiology

## Abstract

K_2P_ potassium channels regulate excitability by affecting cellular resting membrane potential in the brain, cardiovascular system, immune cells, and sensory organs. Despite their important roles in anesthesia, arrhythmia, pain, hypertension, sleep, and migraine, the ability to control K_2P_ function remains limited. Here, we describe a chemogenetic strategy termed CATKLAMP (<u>C</u>ovalent <u>A</u>ctivation of <u>T</u>REK family <u>K</u>^+^ channels to c<u>LA</u>mp <u>M</u>embrane <u>P</u>otential) that leverages the discovery of a site in the K_2P_ modulator pocket that reacts with electrophile-bearing derivatives of a TREK subfamily small molecule activator, ML335, to activate the channel irreversibly. We show that the CATKLAMP strategy can be used to probe fundamental aspects of K_2P_ function, as a switch to silence neuronal firing, and is applicable to all TREK subfamily members. Together, our findings exemplify a new means to alter K_2P_ channel activity that should facilitate studies both molecular and systems level studies of K_2P_ function and enable the search for new K_2P_ modulators.

## Introduction

Leak potassium currents produced by K_2P_ channels play a fundamental role in membrane potential stabilization and cellular excitability regulation^1,2^. Consequently, K_2P_s are important to normal and pathophysiological processes^2,3^ such as action potential propagation^4,5^, pain^6-8^, sleep^9^, intraocular pressure^10^, retinal visual processing^11^, migraine^12^, depression^13^, pulmonary hypertension^14^, and sleep apnea^15^. Because of their strong effects on cellular excitability, K_2P_s are highly regulated by a variety of physiological cues and regulatory pathways^2^. Further, various K_2P_ family members are viewed as good therapeutic targets for pain^3,6,16^, anesthetic responses^3,16,17^, migraine^3,12,16^, and sleep apnea^3,16,18^. Recent structural pharmacology studies have established that K_2P_s have a polysite pharmacology in which distinct binding sites for small molecules are found at every layer of the channel structure^16,18-22^. Nevertheless, K_2P_ pharmacology remains underdeveloped. The vast majority of K_2P_ modulators possess only micromolar half maximal effective concentrations (EC_50_s) and show limited selectivity. Further, even though there is structural characterization of modulators for the Keystone inhibitor site^20^, K_2P_ modulator pocket^21,23^, Fenestration site^19,24^, Vestibule site^18^, and Modulatory lipid site^21-23^, most K_2P_ modulators lack well-defined mechanisms of action or structurally defined binding sites^3,16^. Thus, there remains a need to develop more potent and selective means to affect K_2P_ function.

Chemogenetics, an approach using small molecule ligands to engage an engineered protein of interest to affect selective control, has great potential to probe biological systems^25,26^. Chemogenetic tool development, exemplified by efforts on designed receptors and ion channels, generally starts with engineering a well-characterized small molecule ligand:receptor pair to create uniquely selective partners^27-29^. As such, due to their rich pharmacology, G-protein coupled receptors have been a favored target^27,30^. By contrast, ion channel chemogenetic efforts have been limited to members of the pentameric ligand-gated ion channel class where there are an abundance of well-studied ligands^25,26,28,29^. Because potassium channel activation acts as a powerful inhibitor of electrical activity^31^, channels from this class are particularly attractive targets for protein engineering approaches to control excitability^32-39^. However, due to the lack of selective ligands that could be used as a starting point, there are yet no chemogenetic systems for this ion channel class.

Here, we describe development of a chemogenetic approach using covalent activators of the TREK subfamily of K_2P_ potassium channels, termed CATKLAMP for ‘<u>C</u>ovalent <u>A</u>ctivation of TREK family K^+^ channels to c<u>LA</u>mp <u>M</u>embrane <u>P</u>otential’. Unlike other chemogenetic systems that rely on steric interactions for selectivity^25,26,28^, the CATKLAMP system pairs genetically modified K_2P_ channels with electrophilic ligands that gain selectivity through covalent engagement of an engineered channel subunit. Based on structural studies, we identified Ser131, a K_2P_2.1(TREK-1) residue in the K_2P_ modulator pocket that controls the selectivity filter C-type gate^21,23^, as a covalent ligand engagement site. We used this knowledge together with structural and functional studies to develop a chemogenetic pair consisting of CAT335, a maleimide derivative of the K_2P_ modulator pocket activator ML335, and an engineered K_2P_2.1(TREK-1) variant bearing an S131C modification, termed TREK-1^CG*^. CAT335 selectively and irreversibly activates TREK-1^CG*^ but not wild-type K_2P_2.1(TREK-1). Use of CATKLAMP to study tandemly linked K_2P_2.1(TREK-1) channels containing 0, 1 or 2 reactive TREK-1^CG*^ subunits showed that occupation of one of the two K_2P_ modulator pockets is sufficient to activate the channel and uncovered cooperative effects on channel open probability, reinforcing ideas about factors that control the selectivity filter C-type gate^16,21,23,40-43^. The ability to silence a particular type of neuron in a network has great potential for probing brain function^44^ and presents a task well-suited to K_2P_ channel activation. To design such a chemogenetic tool, we combined the TREK-1^CG*^ background with mutations that suppress K_2P_2.1(TREK-1) basal activity^45^ to create channels having low basal leak currents that hyperpolarize the target cell plasma membrane in response to CAT335 activation and are able to silence hippocampal neuronal firing. We further show that the CATKLAMP approach can be generalized to the other members of the TREK subfamily, K_2P_10.1(TREK-2) and K_2P_4.1(TRAAK)^2,16^. Together our findings demonstrate the potential for using chemogenetic-based tools for probing multiple types of K_2P_s and set a direction for using the CATKLAMP methodology to probe K_2P_ function on levels that span from ion channel biophysics to physiological systems.

## Results

### Covalent modification of the K_2P_2.1 (TREK-1) modulator pocket

In the course of characterizing synthetic derivatives of the TREK family activator ML335^21^, we determined the structure of a previously characterized K_2P_2.1(TREK-1) construct, K_2P_2.1(TREK-1)_cryst21_, complexed with ML336, an ML335 acrylamide derivative (Fig. 1A) at 2.9Å resolution by X-ray crystallography (Figs. 1B and S1A, Table S1). The structure showed ML336 bound to the K_2P_ modulator pocket^16,21^ (Fig. 1B) and surprisingly revealed continuous electron density between the ML336 acrylamide and Ser131 sidechain hydroxyl (Fig. S1A), indicating covalent bond formation between ML336 and the channel. The structure shows that ML336 interacts primarily with K_2P_ modulator pocket residues Phe134, Lys271, and Trp275 (Fig. 1B) in a manner similar to ML335^21^. The new covalent bond results from Michael-type reaction of the ML336 acrylamide with the Ser131 hydroxyl, a residue that donates a hydrogen bond to the ML335 methyl sulfonamide moiety in the ML335 structure^21^. As a result of covalent bond formation, the ML336 upper ring sits ∼1.5Å lower in the K_2P_ modulator pocket relative to ML335, whereas its lower ring is positioned similarly to its non-covalent parent ML335 (Fig. S1B), raising the expectation that ML336 could act as an irreversible, covalent K_2P_2.1(TREK-1) activator.

**Figure 1.**
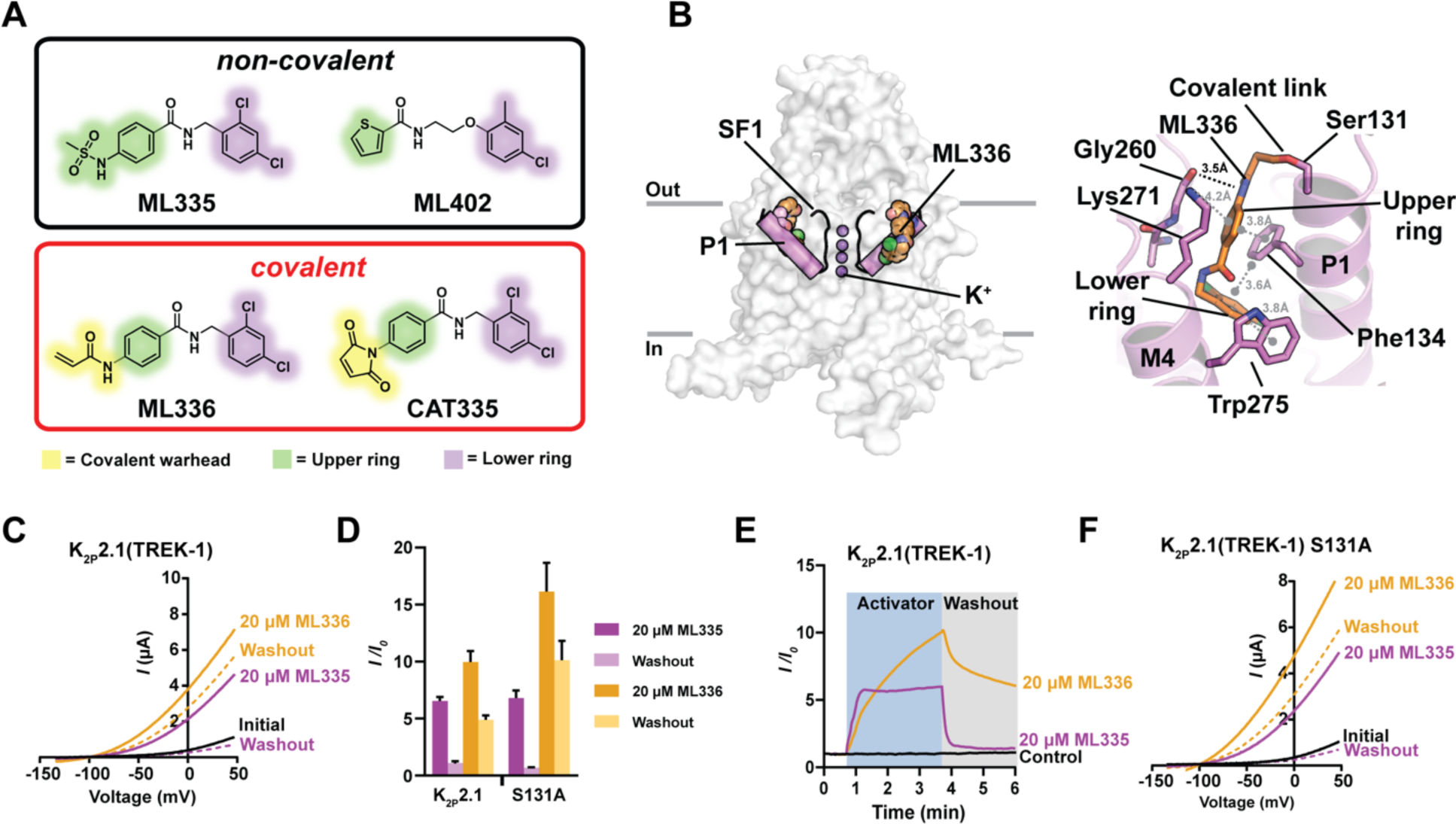
ML336 covalently activates K_2P_2.1(TREK-1) **A,** Chemical structures of non-covalent (top) and covalent (bottom) K_2P_ modulator pocket activators. **B,** (left) K_2P_2.1(TREK-1):ML336 complex structure. P1 and SF1 are highlighted (magenta). Potassium ions (violet) and ML336 (orange) are shown in space filling. Grey lines denote membrane (right) ML336 interaction details. Hydrogen bonds are black. Cation-π and π-π interactions are grey. **C,** Two-electrode voltage clamp (TEVC) traces from *Xenopus* oocytes expressing K_2P_2.1(TREK-1) (black), and activated by 20 µM ML335 (magenta) or 20 µM ML336 (yellow). ‘Washout’ indicates currents recorded following 3 minutes of buffer perfusion for each activator and are color coded. **D,** Activation (*I/I_0_*) of K_2P_2.1(TREK-1) following application and washout of ML335 (magenta) or ML336 (yellow) (n=3-5). **E,** Timecourse of activation (*I/I_0_*) of K_2P_2.1(TREK-1) at 0 mV following addition of either ML335 (magenta) or ML336 (yellow) (blue highlight). Washout for each is shown (grey highlight). Control shows K_2P_2.1 (TREK-1) in the absence of activator. **F,** Exemplar TEVC traces from *Xenopus* oocytes expressing K_2P_2.1(TREK-1) S131A (black) and activated by 20 μM ML335 (magenta) or 20 μM ML336 (yellow). ‘Washout’ indicates currents recorded following 3 minutes of buffer perfusion for each activator and are color coded. Data are mean ± S.E.M.

Based on our crystallographic observations, we used two-electrode voltage clamp (TEVC) in *Xenopus* oocytes to search for functional evidence in support of ML336 irreversible covalent activation of the channel. Comparison of the effects of 20 µM ML335 and ML336 on oocytes expressing K_2P_2.1(TREK-1) showed that ML336 activates K_2P_2.1(TREK-1) (*I*_ML336_/*I*_0_ = 10 ± 1.0) better than ML335 (Figs. 1C-D, Table S2). Moreover, unlike ML335, ML336 activation was sustained following washout over the course of several minutes (Fig. 1E). To determine if this activation was related to covalent adduct formation, we examined the effects of ML335 and ML336 on two K_2P_2.1(TREK-1) mutants: S131A a mutant incapable of forming the covalent link and S131C a mutant that should have enhanced reactivity to the acrylamide. Surprisingly, application of ML336 to K_2P_2.1(TREK-1) S131A activated the channel similar to K_2P_2.1(TREK-1) (Figs. 1C, D, and F, Table S2). Further, although 5 µM ML336 application to K_2P_2.1(TREK-1) S131C (denoted as TREK-1^CG*^) activated the channel, the compound showed washout behavior similar to the ML336:K_2P_2.1(TREK-1) combination, contrary to the expected reaction enhancement from the cysteine substitution (Figs. S1C-E). Thus, even though ML336 could form a covalent adduct with the K_2P_2.1(TREK-1) Ser131 under crystallization conditions (high concentrations of both components over a week long timeframe) we found no evidence for covalent adduct formation over the minute timescales used for functional studies. Nevertheless, discovery of the ML336-Ser131 adduct revealed the availability of residue 131 to be modified covalently by K_2P_ modulator pocket activators and prompted us to explore alternative approaches to create a covalent K_2P_2.1(TREK-1) modulator.

### Development of an activator:K_2P_2.1(TREK-1) chemogenetic pair

To develop a small molecule:K_2P_ pair possessing rapid, irreversible activation we explored the consequences of incorporating other electrophilic functional groups in place of the ML336 acrylamide. We identified the maleimide CAT335 for ‘<u>C</u>ovalent <u>A</u>ctivator of <u>T</u>REK, ML<u>335</u> scaffold’ as a suitable compound (Fig. 1A, Supplementary Scheme 1) and paired this activator with the TREK-1^CG*^ target channel bearing a cysteine at the site expected to react with the CAT335 maleimide (S131C). Biophysical studies showed that the S131C change did not affect the EC_50_ for the non-covalent agonist ML335 (Fig. S2A) (EC_50_ = 18 ± 4 μM and 24 ± 4 μM; for TREK-1^CG*^ and K_2P_2.1(TREK-1) respectively) or activation by temperature (Fig. S2B) relative to wild-type K_2P_2.1(TREK-1). However, we did observe a reduced efficacy of ML335 activation (E_max_ *I*_ML335_/*I*_0_ = 9.6 ± 0.8 and 15.8 ± 1.3, for TREK-1^CG*^ and K_2P_2.1(TREK-1) respectively) (Fig. S2A, Table S2) and a change in the extracellular pH response (Fig. S2C) indicating that the S131C change affected some channel properties, possibly due to the proximity of S131C to His126, a key residue for K_2P_2.1(TREK-1) extracellular pH sensing^46^. Nevertheless, these data indicate that TREK-1^CG*^ largely functions similarly to the parent channel.

Application of 20 µM CAT335 to *Xenopus* oocytes expressing TREK-1^CG*^ strongly activated the channel but by contrast evoked only minimal response from K_2P_2.1(TREK-1) (*I*/*I*_0_ = 22 ± 2 and 2.0 ± 0.2 for TREK-1^CG*^ and K_2P_2.1(TREK-1), respectively) (Figs. 2A-C, Table 1). Comparison of washout time courses for TREK-1^CG*^ (Fig. 2D) revealed an immediate reversal of the effects of the non-covalent activator ML335, matching its behavior on K_2P_2.1(TREK-1) (Fig. S2D) that contrasted the persistent activation of TREK-1^CG*^ by CAT335. K_2P_2.1(TREK-1) showed some residual activation of by 20 µM CAT335 following washout (*I*_CAT335_/*I*_0_ = 4.0 ± 0.6 after washout) (Figs. 2B and S2D). This latent effect was absent at 5 µM CAT335, a concentration that potently activated TREK-1^CG*^ (Figs. S2E-F). To examine whether this latent K_2P_2.1(TREK-1) activation was due to the use of oocytes, we used whole cell patch clamp electrophysiology to characterize the CAT335 responses of TREK-1^CG*^ and K_2P_2.1(TREK-1) in HEK293 cells. As in oocytes, the non-covalent activator ML335 activated both channels but was slightly less effective at stimulating TREK-1^CG*^ (*I*_ML335_/*I*_0_ = 3.7 ± 0.4 and 6.2 ± 0.5 at 20µM, TREK-1^CG*^ and K_2P_2.1(TREK-1), respectively (Table 1). Further, CAT335 caused a strong, irreversible activation of TREK-1^CG*^, but elicited a modest, reversible response from K_2P_2.1(TREK-1) (*I*_CAT335_/*I*_0_ = 8.5 ± 1.5 and 1.9 ± 0.2 for TREK-1^CG*^ and K_2P_2.1(TREK-1), respectively) (Figs. 2E-H, Table 1). Hence, the observed latent activation in oocytes was a consequence of the expression system and not a general CAT335 property.

**Figure 2.**
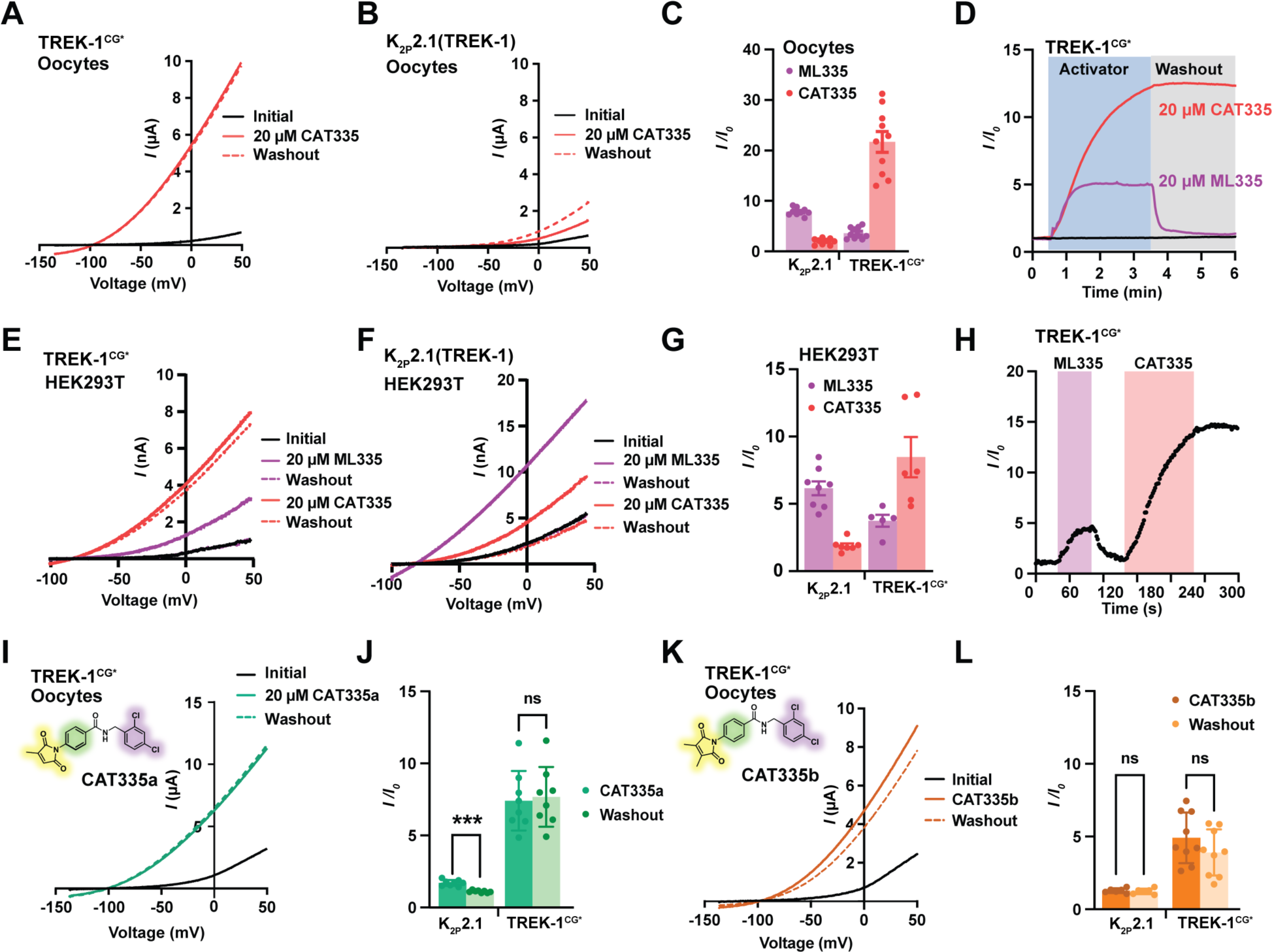
Properties of TREK-1^CG*^:covalent ligand pairs. **A-B,** Exemplar TEVC responses of *Xenopus* oocytes expressing **A,** TREK-1^CG*^ or **B,** K_2P_2.1(TREK-1) to application of 20 µM non-covalent (ML335) (magenta) and covalent (CAT335) (red) activators. **C,** Activation (*I/I_0_*) and washout response of oocytes expressing K_2P_2.1(TREK-1) or TREK-1^CG*^ to application of 20 μM ML335 (magenta) or 20 μM CAT335 (red) (n=9-10). **D,** Exemplar TEVC time courses of TREK-1^CG*^ activation by 20 μM ML335 (magenta) and 20 μM CAT335 (red). **E-F,** Exemplar whole-cell currents from HEK293 cells expressing **E,** TREK-1^CG*^ or **F,** K_2P_2.1(TREK-1) to application of 20 µM non-covalent (ML335) (magenta) and covalent (CAT335) (red) activators. **G,** Activation (*I/I_0_*) of HEK293 cells expressing K_2P_2.1(TREK-1) or TREK-1^CG*^ following application of 20 μM ML335 (magenta) or 20 μM CAT335 (red) (n=5-8). **H,** Exemplar time course of TREK-1^CG*^ activation in HEK293 cells by 20 μM ML335 (magenta) and 20 μM CAT335 (red). **I,** Exemplar TEVC responses of TREK-1^CG*^ to 20 μM CAT335a (green). **J,** Activation (*I/I_0_*) and washout response of oocytes expressing TREK-1^CG*^ to 20 μM CAT335a (green) (n=8). **K,** Exemplar TEVC responses of TREK-1^CG*^ to 20 μM CAT335b (orange). **L,** Activation (*I/I_0_*) and washout response of oocytes expressing TREK-1^CG*^ to 20 μM CAT335b (orange) (n=6-9). Significance was measured for panels J and L using unpaired t-tests (n.s. p>0.12; *** p<0.0002). Error bars represent S.E.M..

**Table 1:**
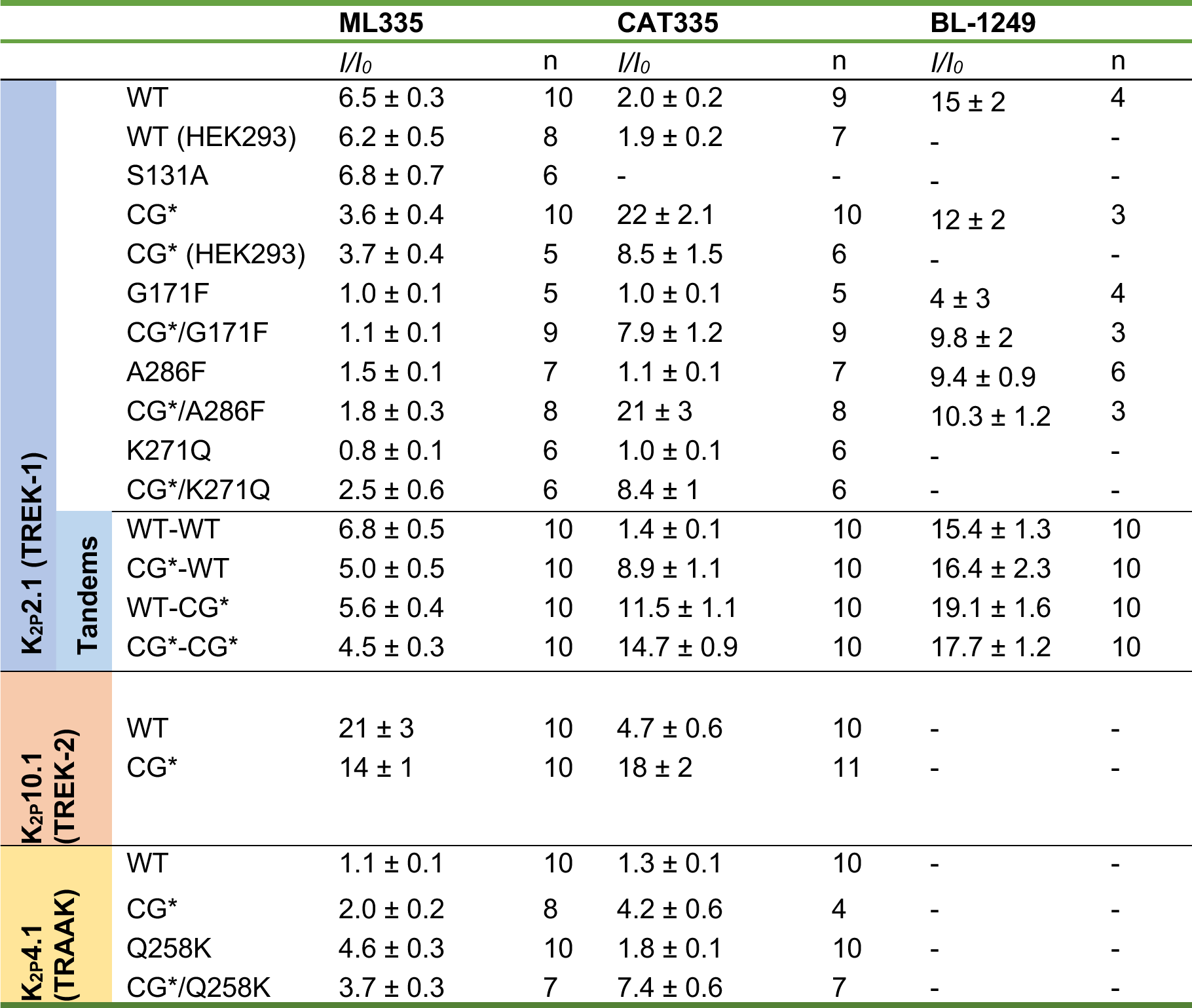
Responses to 20 µM activator.

To explore the structure-function relationship between CAT335 and the K_2P_ modulator pocket, we synthesized a set of CAT335 derivatives bearing one or two methyl groups on the reactive maleimide moiety (CAT335a and CAT335b) and a derivative with an isosteric but non-electrophilic succinimide ring (CAT335c) (Supplementary Scheme 1). The methyl-substituted maleimide CAT335a irreversibly activated TREK-1^CG*^ but had a reduced response magnitude relative to CAT335 (*I*_CAT335a_/*I*_0_ = 7.4 ± 0.7 at 20 µM) (Figs. 2A and I, C and J, Table S2). The dimethyl-substituted CAT335b was also an irreversible TREK-1^CG*^ activator but was not as effective as CAT335 or CAT335a (*I*_CAT335b_/*I*_0_ = 4.9 ± 0.6 at 20 µM) (Figs. 2K-L, Table S2). The non-electrophilic CAT335c reversibly activated TREK-1^CG*^ and weakly activated K_2P_2.1(TREK-1) (*I*_CAT335c_/*I*_0_ = 5.0 ± 0.4 and 2.1 ± 0.1 at 20 µM for TREK-1^CG*^ and K_2P_2.1(TREK-1), respectively) (Figs. S3A-C. Table S2). Hence, none of the modifications improved behavior relative to the starting CAT335 molecule.

To test whether the activation differences among the three maleimide-based activators CAT335, CAT335a, and CAT335b reflected differences in channel covalent modification, we applied each at concentrations spanning 0-20 µM for 1 hour to oocytes expressing TREK-1^CG*^ or K_2P_2.1(TREK-1). We then measured the resultant basal currents, assessed whether there were unmodified remaining channels that could be activated by 20 µM ML335, and determined the extent to which the cumulative activation was reversible. These experiments showed that extended incubation times increased the extent of activation evoked by all three maleimides with a rank order of CAT335>CAT335a>>CAT335b (Figs. S3D-F). Although CAT335 and CAT335a showed some remaining ML335-activatable current at 1 µM (∼20%), this residual, activatable current was absent when either compound was applied at concentrations ≥5 µM (Figs.S3D-E). Notably, at CAT335 and CAT335a concentrations ≥5 µM activation was irreversible, indicating that both compounds had completely labeled the available TREK-1^CG*^ channels. By contrast, for CAT335b the residual ML335 activation only disappeared at 20 µM and activation remained reversible at all tested concentrations. These differences from CAT335 and CAT335a indicate that the introduction of methyl groups at both electrophilic carbon atoms of the maleimide caused steric occlusion that interfered with the ability of CAT335b to bind and activate the channel irreversibly. Importantly, 20 µM application of each of the covalent activators for 1 hour had no effect on K_2P_2.1(TREK-1) (Figs. S3D-F), establishing that the irreversible activation effects were a consequence of the malimide:S131C combination.

To probe the differences in reactivity between CAT335 and CAT335a further, we examined the kinetics of TREK-1^CG*^ engagement in *Xenopus* oocytes to calculate both the affinity (K_a_) and reactivity components (k_act_) of the interaction (Fig. S4). These studies revealed that CAT335 is more potent than CAT335a by ∼3-fold (k_act_/K_a_ = 1.5x10^-3^ M^-1^min^-1^ vs. k_act_/K_a_ = 0.5x10^-3^ M^-1^min^-1^, respectively), and that the higher CAT335 potency is driven by reactivity differences between the CAT335 and CAT335a (k_act_=.068 min^-1^ vs. k_act_=.017 min^-1^, respectively) rather than affinity differences (K_a_ = 46.87 μM vs. K_a_ = 35.54 μM, respectively) (Figs. S4C-D). Because the persistent activation of K_2P_s is expected to stabilize the target cell membrane potential at the potassium reversal potential, we named this chemogenetic pair system comprising the maleimide-bearing activator and target TREK-1^CG*^ channel as ‘CATKLAMP’ for <u>C</u>ovalent <u>A</u>ctivator of <u>T</u>REK <u>K</u>^+^ channels whose action c<u>LA</u>mps the <u>M</u>embrane <u>P</u>otential near the potassium reversal potential.

### CATKLAMP structures show a binding mode matching ML335

To understand the interactions of the CATKLAMP partners, we crystallized and determined the structures of TREK-1^CG*^ complexed with CAT335, CAT335a, and the non-covalent activator ML335 at resolutions of 3.0Å each (Figs. 3A-D, S5A-C and Table S1). These structures showed no major global deviations from the K_2P_2.1(TREK-1):ML335 complex^21^ (Figs. S5D) (Root mean square deviation (RMSD)Cα = 0.446Å, 0.554Å, and 0.335Å for CAT335, CAT335a, and ML335 complexes versus PDB:6CQ8). Importantly, both the CAT335 and CAT335a structures showed continuous density that bridged the maleimide moiety and S131C, indicative of covalent adduct formation (Figs. 3A-C, S5A-B) and supporting the functional observations showing that both compounds act on TREK-1^CG*^ in an irreversible manner (Figs. 2A, D, E, H, I, J, and S2F). The K_2P_ modulator pocket residues that surround the CAT335 and CAT335a (Phe134, Gly260, Lys271, and Trp275) make interactions with both compounds that are similar to those seen in the K_2P_2.1(TREK-1):ML335 structure (Fig. 3A-C) and in the non-covalent TREK-1^CG*^:ML335 complex (Fig. 3D-E). Further, the CAT335 and CAT335a upper and lower ring positions largely match those of the non-covalent ML335 complex (Fig. 3C). Thus, the structures show that the CAT335 ligands make a covalent bond with the S131C target similar to the acrylamide-bound K_2P_2.1(TREK-1):ML336 covalent adduct (Fig. 1B) that inspired their design and use the ML335 binding mode to engage the K_2P_ modulator pocket. Hence, formation of the covalent bond enhances binding by irreversibly trapping the ML335 core scaffold in the K_2P_ modulator pocket.

**Figure 3.**
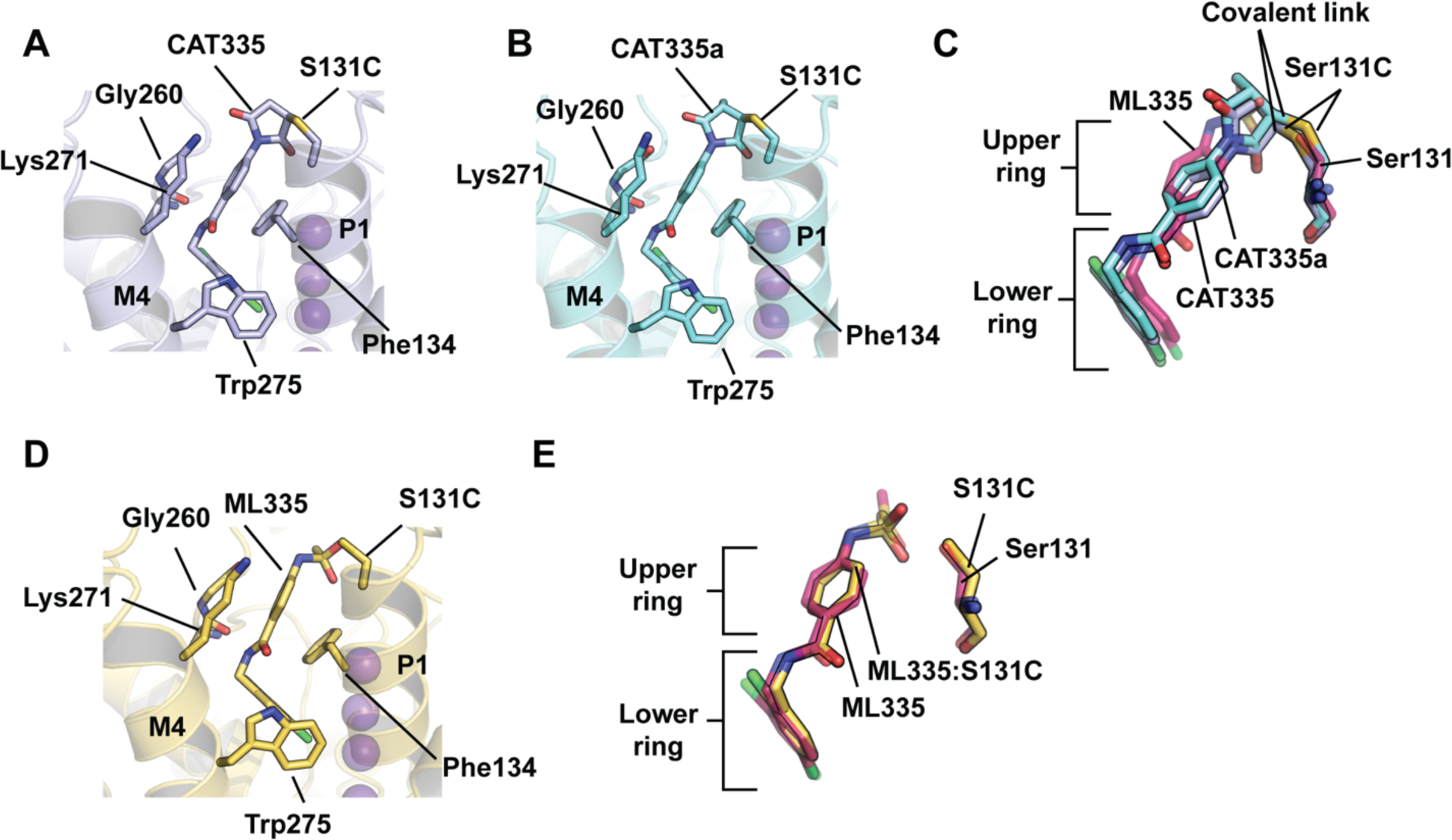
TREK-1^CG*^ ligand interactions. **A-B,** Cartoon diagram of **A,** TREK-1^CG*^:CAT335 (light blue) and **B,** TREK-1^CG*^:CAT335a (aquamarine) K_2P_ modulator pocket interactions. **C,** Superposition of CAT335 (light blue) and CAT335a (aquamarine) with ML335 (warm pink) from the K_2P_2.1 (TREK-1):ML335 complex (PDB:6CQ8)^21^. Residue 131, covalent links, and upper and lower rings are indicated. **D,** Cartoon diagram of TREK-1^CG*^:ML335 (yellow) interactions. **E,** Superposition of ML335 from the TREK-1^CG*^:ML335 (yellow) and K_2P_2.1 (TREK-1):ML335 (PDB:6CQ8)^21^ (warm pink) complexes. Residue 131 and upper and lower rings are indicated. Potassium ions in ‘A’, ‘B’, and ‘D’ are shown as purple spheres.

### Concatenated Tandem Channels reveal K_2P_ modulator pocket cooperativity

ML335 binding to the K_2P_2.1(TREK-1) K_2P_ modulator pocket provides a profound stabilization to the selectivity filter C-type gate that is the principal control point for most K_2P_s^16,23,40,41^. However, it has been unclear whether occupation of one or both K_2P_ modulator pocket sites present on the channel is required to stabilize the selectivity filter and activate the channel. Given the ability of CAT335 to activate TREK-1^CG*^ selectively relative to K_2P_2.1(TREK-1) (Figs. 2C and G), we created a set of tandem channels, following our previous tandem K_2P_ designs^47^ in which the two K_2P_ dimer subunits were covalently concatenated, bearing zero (WT-WT), one (CG*-WT and WT-CG*), or two (CG*-CG*) TREK-1^CG*^ subunits and used these to test the consequences of activating a single subunit at the K_2P_ modulator pocket (Fig. 4A). To probe the responses of the channels to both non-covalent and covalent activators, we performed an experiment in which we tested the effects of sequential activation by ML335, CAT335, and an activator that stabilizes the selectivity filter by acting on a separate part of the channel, BL-1249^16,19,48^ (Fig. 4A). Measurement of activation by 20 µM ML335 showed that both homomeric constructs responded similarly to their unconcatenated counterparts (*I*_ML335_/*I*_0_ = 6.8 ± 0.5 and 4.5 ± 0.3 for WT-WT and CG*-CG*, respectively) (Figs. 4A-B and S2A, Table 1). Moreover, the response of the tethered heterodimers was similar to each other (*I*_ML335_/*I*_0_ = 5.0 ± 0.5 and 5.6 ± 0.4 for CG*-WT and WT-CG*, respectively) (Fig. 4A-B). Together, these results indicate that the tandem linkage has minimal effect on K_2P_ modulator pocket activation, in line with its absence of effects on other gating modalities^47^.

**Figure 4.**
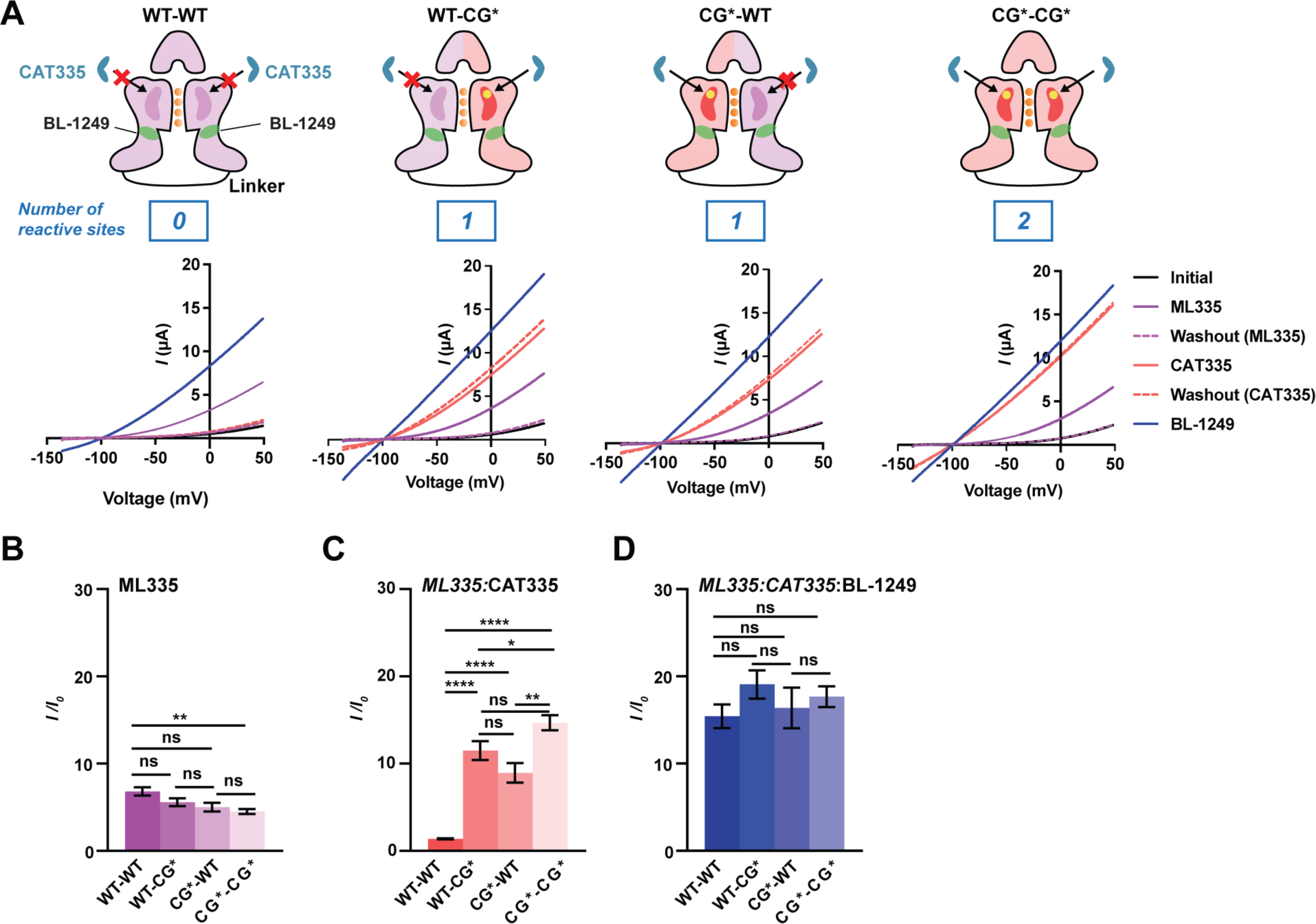
CAT335 responses of tandem K_2P_s having varied numbers of reactive sites. **A,** Exemplar TEVC traces from *Xenopus* oocytes expressing tandem WT-WT, WT-CG*, CG*-WT, and CG*-CG* K_2P_2.1(TREK-1) channels following sequential addition and washout of 20 μM ML335 (magenta), 20 μM CAT335 (red), and 20 μM BL-1249 (blue). Cartoons show tandem construct schematics. Number of CAT335 reactive sites per channel and BL-1249 binding sites (green) are indicated. **B-D,** Tandem K_2P_2.1(TREK-1) channel activation (*I/I_0_*) following sequential application of **B,** 20 μM ML335 (magenta), **C,** 20 μM CAT335 (red) after 20 μM ML335 and **D,** 20 μM BL-1249 (blue) after 20 μM ML335 and 20 μM CAT335 (n=10). Prior agonists are shown in italics. Significance was measured using Brown-Forsythe and Welch one-way ANOVA with Dunnett’s T3 multiple comparisons test (n.s. p>.1234; * p<.0332; **p<.0021; *** p<.0002; **** p<.0001). Error bars are S.E.M.

We next tested the sensitivity of the homodimeric and heterodimeric channels to the covalent activator CAT335. As expected, application of 20 μM CAT335 had little effect on WT-WT but caused strong, irreversible activation of CG*-CG*, recapitulating the results obtained with unlinked homomeric channels (*I*_CAT335_/*I*_0l_ = 1.4 ± 0.1 and 14.7 ± 0.9 for WT-WT and CG*-CG*, respectively) (Figs. 4A and C, Table 1). Notably, CAT335 was able to activate both heterodimeric channels WT-CG* and CG*-WT to a similar degree (*I*_CAT335_/*I*_0_= 11.5 ± 1.1 and 8.9 ± 1.1, respectively), establishing that occupation of a single K_2P_ modulator pocket is sufficient increase channel function. These responses represent 60-80% of CG*-CG* homodimer activation (Fig. 4A and C, Table 1). The application of 20 µM BL-1249 after ML335 and CAT335 resulted in a similar degree of total activation for all constructs (Figs. 4A and D, Table 1). Compared to WT-WT, BL-1249 had less stimulatory effect on CAT335 treated WT-CG* and CG*-WT channels (Fig. 4A), and almost no stimulatory effect on CAT335 treated CG*-CG* channels, indicating that CAT335 covalent-mediated activation of the C-type selectivity filter gate reduces the response of TREK-1^CG*^-containing constructs to BL-1249.

To understand the nature of CAT335 activation further, we turned to single channel recording. Because WT-CG* and CG*-WT had similar responses to CAT335 (Fig. 4, Table 1), we focused on CG*-WT and compared its behavior with WT-WT and CG*-CG* channels. Single channel current amplitudes were similar for all three constructs (Figs. 5A-C, S6A-B). WT-WT and CG*-WT had similar basal open probabilities (P_o_), but this value was ∼2-fold larger for CG*-CG* (P_o basal_ = 0.020, 0.027, and 0.058 for WT-WT, CG*-WT, and CG*-CG*, respectively) (Fig. S6C), highlighting the sensitivity of the K_2P_ modulator pocket to physicochemical perturbations such as the S131C change. As in the whole cell experiments, 20 µM CAT335 had no impact on WT-WT. By contrast, 20 µM CAT335 caused large increases in P_o_ when applied to either construct containing its chemogenetic target subunit (P_o CAT335_/P_o basal_ = 6.3 and 7.8 for CG*-WT and CG*-CG*, respectively) (Figs. 5D-F). Recapitulating the effects of ML335 on K_2P_2.1(TREK-1)^23^, CAT335 elicited a large P_o_ increase for all CG* tandem channels but had minimal effect on single channel conductance at -100 mV and +50 mV (Figs. 5J-O). By contrast, application of 20 µM BL-1249 alone resulted in an increase in both single channel conductance at -100 mV and P_o_ for all three tandem channels (Figs. 5G-I and S6D-F), consistent with its reported effects on K_2P_s (Fig. S6G)^19^. Hence, the activation differences observed for CAT335 are a direct consequence of the engagement of this molecule with the target subunits. Taken together with the structural data (Figs. 3A and C), these results indicate that like its non-covalent parent, ML335^23^, CAT335 activation results from conductive state stabilization through its action on the K_2P_ modulator pocket.

**Figure 5.**
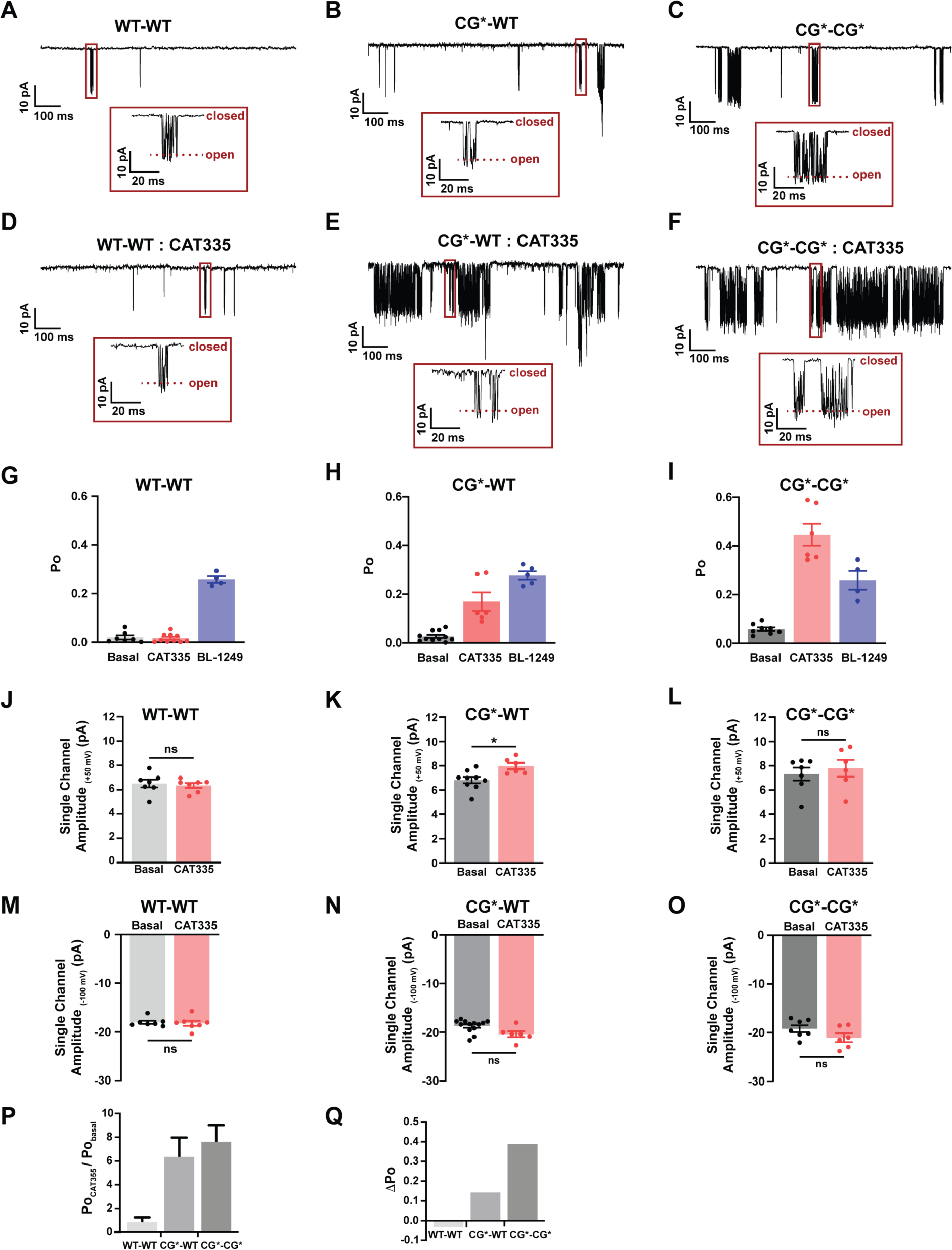
Single channel analysis of K_2P_2.1(TREK-1) tandems. **A-C,** Exemplar cell attached single channel recordings at -100 mV from **A,** WT-WT, **B,** CG*-WT, and **C,** CG*-CG*. **D-F,** Exemplar cell attached single channel recordings from **D,** WT-WT, **E,** CG*-WT, and **F,** CG*-CG* after treatment with 20 μM CAT335. **G-I,** Open probability (P_o_) of **G,** WT-WT, **H,** CG*-WT, and **I,** CG*-CG* under basal conditions (gray), after application of 20 μM CAT335 (red), or after application of 20 μM BL-1249 (blue) (n=4-11). Insets in A-F show boxed regions. **J-L,** Single channel amplitude at +50 mV for: **J,** WT-WT, **K,** CG*-WT, and **L,** CG*-CG* channels under basal conditions (gray) or with 20 μM CAT335 (red) (n=6-13). **M-O,** Single channel amplitude at -100 mV for: **M,** WT-WT, **N,** CG*-WT, and **O,** CG*-CG* channels under basal conditions (gray) or with 20 μM CAT335 (red). **P,** P_o_ ratios for the indicated constructs. **Q,** Λ1P_o_ (P_o CAT335_-P_o control_) for the indicated constructs. Significance was measured using Brown-Forsythe and Welch one-way ANOVA with Dunnett’s T3 multiple comparisons test (n.s. p>0.12; * p<0.033; **p<0.0021; *** p<0.0002; **** p<0.0001). Error bars are S.E.M.

The active conformation of the K_2P_2.1(TREK-1) selectivity filter is influenced by the presence of permeant ions and K_2P_ modulator pocket occupation^23^. The selectivity of our chemogenetic approach afforded the opportunity to engage either a one, CG*-WT, or both, CG*-CG*, K_2P_ modulator pockets to assess how the stoichiometry of activator binding affects channel activation. As expected, the fractional change in P_o_ (P_o CAT335_/P_o Basal_) (Fig. 5P) paralleled that seen in the TEVC experiments (Fig. 4C) with CG*-WT showing ∼80% of the fractional activation of CG*-CG*. Notably, the change in P_o_ (ΔP_o_) caused by CAT335 was ∼3-fold larger in the tandem channel having two reactive sites (Fig. 5Q). This non-additive P_o_ change between CG*-CG* and CG*-WT indicates that the two modulator pocket sites are coupled. Observation of this coupling corroborates the idea that the selectivity filter conformation and its ability to coordinate permeating ions depends on the conformation of the surrounding channel structure^21^.

### Attenuated TREK-1^CG*^ mutants create a switch-like chemogenetic pair and highlight Modulator pocket and Fenestration site coupling

Because of their intrinsic basal activity, expression of K_2P_2.1(TREK-1)^32,49^ or TREK-1^CG*^ can hyperpolarize the target cell. We sought to reduce this basal activity while retaining the TREK-1^CG*^ CAT335 response. Previous studies have shown that G171F and A286F mutations on the pore lining M2 and M4 helices, respectively, yield channels having a low basal current that can be activated by flufenamic acid (FFA)^45^. This compound is a member of the fenamate class of activators that includes BL-1249 that act at a lateral gap below the selectivity filter lined by the M2 and M4 helices known as the ‘fenestration site’ (FS)^16,19,24^ (Fig. 6A). As the M2 and M4 transmembrane helices are coupled to the selectivity filter gate^40,47,50,51^, we tested whether G171F and A286F mutations could create a switch-like chemogenetic pair in which an ‘off’ basal state could be switched irreversibly to ‘on’ by CAT335.

**Figure 6:**
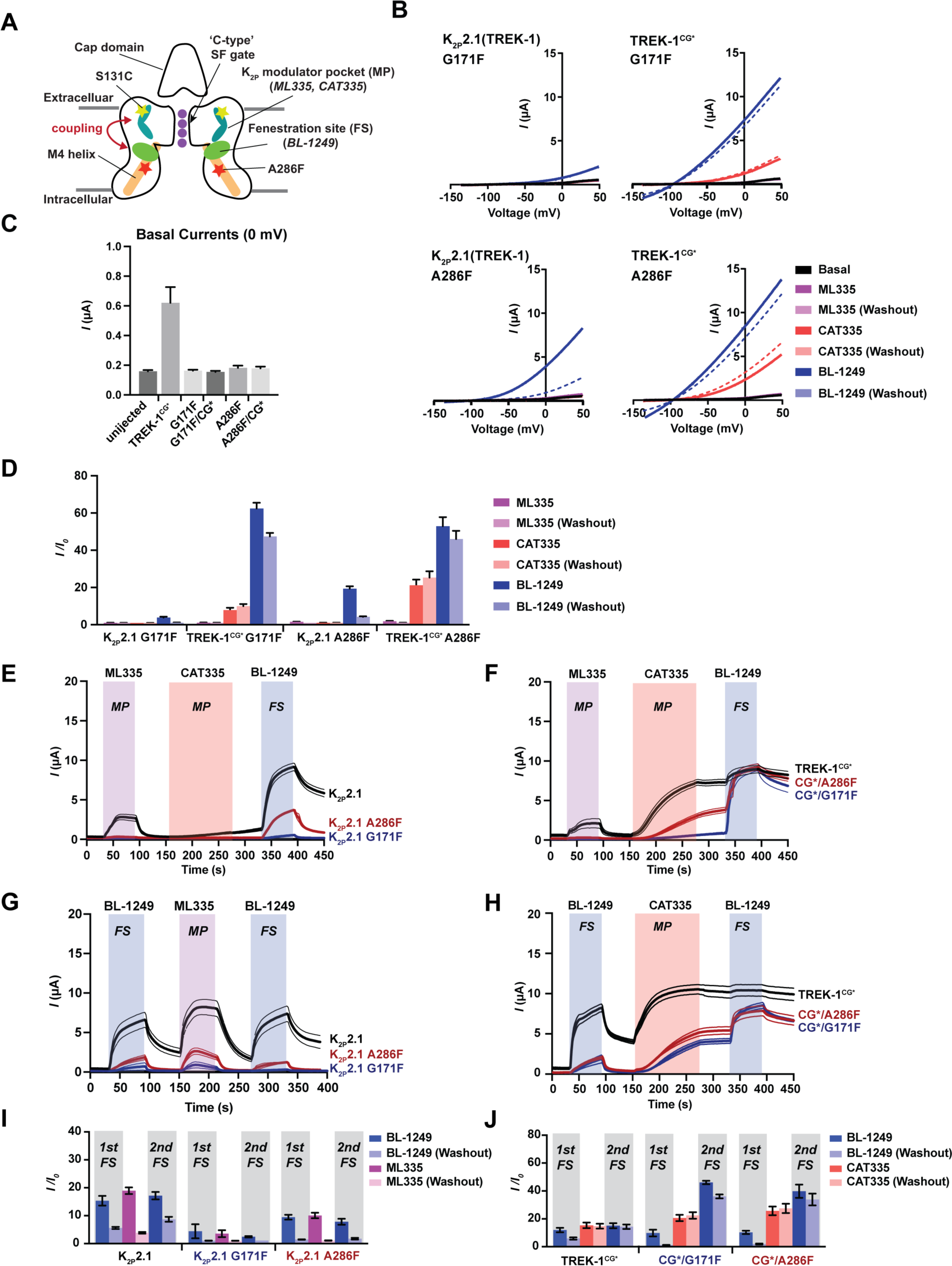
Pharmacological responses of attenuated TREK-1^CG*^ mutants. **A,** Cartoon diagram showing locations of the K_2P_ modulator pocket (MP) (cyan), Fenestration site (FS) (green), S131C, and A286F. G171F is on a similar level as A286F. Select channel elements and coupling between MP and FS sites are labeled. **B,** Exemplar TEVC current traces for K_2P_2.1(TREK-1) G171F, K_2P_2.1(TREK-1) A286F, TREK-1^CG*^ G171F, and TREK-1^CG*^ A286F following sequential application of 20 μM ML335 (magenta), 20 μM CAT335 (red), and 20 μM BL-1249 (blue). **C,** Basal currents at 0 mV for the indicated channels in *Xenopus o*ocytes. **D,** Activation (*I/I_0_*) for the indicated channels at 0 mV in response to the sequential addition of 20 μM ML335 (magenta), 20 μM CAT335 (red), and 20 μM BL-1249 (blue) (n=5-8). **E-H,** Timecourses of K_2P_ modulator pocket (MP) and Fenestration site (FS) activator responses at 0 mV. **E-F,** Responses to application of 20 μM ML335 (magenta), 20 μM CAT335 (red), and 20 μM BL-1249 (blue) during the highlighted windows for **E,** K_2P_2.1(TREK-1) (black), K_2P_2.1(TREK-1) G171F (blue), and K_2P_2.1(TREK-1) A286F (red) and **F,** TREK-1^CG*^ (black), TREK-1^CG*^ G171F (blue), and TREK-1^CG*^A286F (red). **G,** K_2P_2.1(TREK-1) (black), K_2P_2.1(TREK-1) G171F (blue), and K_2P_2.1(TREK-1) A286F (red) responses to application of 20 μM BL-1249 (blue), 20 μM ML335 (magenta) or 20 μM CAT335 (red), and 20 μM BL-1249 (blue) during the highlighted windows. **H,** TREK-1^CG*^ (black), TREK-1^CG*^ G171F (blue), and TREK-1^CG*^A286F (red) responses to application of 20 μM BL-1249 (blue), 20 μM ML335 (magenta) or 20 μM CAT335 (red), and 20 μM BL-1249 (blue) during the highlighted windows. For **E-H** thick lines represent mean values, thin lines represent S.E.M. (n=3-6). **I,** Activation (*I/I_0_*) for the indicated channels at 0 mV in response to the sequential addition of BL-1249 (blue) 20 μM ML335 (magenta), and 20 μM BL-1249 (blue) (n=5-8)**, J,** Activation (*I/I_0_*) for the indicated channels at 0 mV in response to the sequential addition of BL-1249 (blue) 20 μM CAT335 (red), and 20 μM BL-1249 (blue) (n=5-8). Error bars are mean ± S.E.M.

Expression of K_2P_2.1(TREK-1) and TREK-1^CG*^ bearing either G171F or A286F mutations revealed that all four channels displayed TEVC basal currents that were not different from control oocytes (Figs. 6B-C and S7A-B), recapitulating the strong basal current suppression reported for these mutants in mammalian cells^45^. Both mutations are outside of the K_2P_ modulator binding pocket (MP)^21,23^ (Fig. 6A). Nevertheless, serial application of ML335, CAT335, and BL-1249 showed that G171F and A286F greatly reduced K_2P_2.1(TREK-1) and TREK-1^CG*^ sensitivity to activation by 20 μM ML335 (*I*_ML335_/*I*_0_ ∼1-1.8, Table 1) (Fig. 6B, D-F, *cf*. Fig. 2C). Consistent with the absence of S131C, K_2P_2.1(TREK-1) G171F and K_2P_2.1(TREK-1) A286F were insensitive to 20 μM CAT335 (Figs. 6B, D, and E, Table 1). By contrast, despite their ML335 insensitivity both TREK-1^CG*^ G171F and TREK-1^CG*^ A286F were activated by 20 μM CAT335, (*I*_CAT335_/*I*_0_ = 7.9± 1.2 and 21 ± 3 for TREK-1^CG*^ G171F and TREK-1^CG*^ A286F, respectively) (Figs. 6B, D-F, Table 1), although the responses were slower and smaller than for TREK-1^CG*^ (Fig. 6F).

These experiments also revealed diminished responses in the G171F and A286F mutants to the FS activator BL-1249 relative to its strong (∼15 fold) activation of K_2P_2.1(TREK-1)^48^. BL-1249 was a particularly ineffective agonist for K_2P_2.1(TREK-1) G171F (*I*_BL-1249_/*I*_0_ = 4.4 ± 3.0 at 20 µM) (Figs. 6B-E, Table 1). However, this compound strongly activated TREK-1^CG*^ G171F (*I*_BL-1249_/*I*_0_ = 62 ± 3) following treatment of this channel with CAT335 (Figs. 6B-D, F). TREK-1^CG*^ A286F was responsive to BL-1249 (*I*_BL-1249_/*I*_Control_ = 10.3 ± 1.2) but also showed substantial sensitization to BL-1249 (*I*_BL-1249_/*I*_0_ = 53 ± 5) after activation by CAT335 (Figs. 6B, D, and F, Table S3). Both post-CAT335 responses were very different from that observed for TREK-1^CG*^, which could not be activated further by BL-1249 following CAT335 covalent activation (Figs. 4A, 6F, Table S3). The observation that BL-1249 responses appear dependent on prior exposure to the covalent activator suggest that the MP and FS binding sites act in a coupled manner in these constructs having attenuated basal activity.

To examine these effects further, we changed the order in which we applied MP and FS activators (Figs. 6G-J). These experiments showed that the MP site activator ML335 had no effect on K_2P_2.1(TREK-1) G171F or K_2P_2.1(TREK-1) A286F when this compound was applied first (Figs. 6B, D and E). By contrast, activation by 20 μM BL-1249 sensitized both channels to subsequent MP activator application (Figs. 6G and I) causing clear changes in the rate and extent of CAT335 activation, indicating that BL-1249 activation primed the MP response. A similar BL-1249 ‘priming’ effect was observed for CAT335 (Figs. 6E and H). This was most pronounced for TREK-1^CG*^ G171F where FS activator simulation enhanced the CAT335 response from *I*_CAT335_/*I*_0_ = 7.9 ± 1.5 to *I*_CAT335_/*I*_0_ = 21 ± 2 (Figs. 6D and H). Hence, these data indicate that there is strong synergistic coupling between the MP and FS sites.

We next examined the effect of simultaneous application of BL-1249 with ML335 or CAT335 (Fig. S7). Simultaneous addition of 10 μM BL-1249 and 10 μM ML335 activated K_2P_2.1(TREK-1) dramatically more than application of 20 μM ML335 or 20 μM BL-1249 alone (*I*_max_/*I*_0_ = 38 ± 3, 6.5 ± 0.3, and 15 ± 2, respectively) (Tables 1 and S3) and produced similarly larger responses in K_2P_2.1(TREK-1) A286F (*I*_max_/*I*_0_ = 22 ± 4, 1.5 ± 0., and 10.3 ± 1.2, respectively). These effects were less apparent in K_2P_2.1(TREK-1) G171F (*I*_max_/*I*_0_ = 4.7 ± 2, 1.0 ± 0.1, and 4 ± 3, respectively) (Figs. S7A and C, Tables 1 and S3). The synergistic effect was even stronger for CG* targets treated with BL-1249 and CAT335. TREK-1^CG*^ quickly reached maximal activation upon 10 µM simultaneous application of both compounds (*I*_max_/*I*_Control_ = 25 ± 7). Even more striking, this concentration of both activators caused substantial activation of TREK-1 CG* G171F and TREK-1 CG*A286F (*I*_max_/*I*_0_ = 53.9 ± 1.5 and *I*_max_/*I*_0_ = 70 ± 3, respectively) (Figs. S7B and D, Table S3). The synergy between BL-1249, which binds in the FS opened by M4 transmembrane helix movement^16,19^, and K_2P_ modulator pocket activators that bind behind the selectivity filter^16,21,23^, indicates that the two pharmacological sites are coupled (Fig. 6H) and corroborate the idea that M4 helix movement affects the conformation of the selectivity filter and its surrounding structure_21,23,40,47,48_.

### CATKLAMP stabilizes resting membrane potential and silences neuronal firing

K_2P_ activation should strongly modulate resting membrane potential (RMP)^2,3,16^. Hence, we explored the ability of TREK-1^CG*^ and TREK-1^CG*^ A286F to affect RMP in HEK293 cells. TREK-1^CG*^ expression resulted in large basal currents (660 ± 110 pA at 0 mV) that hyperpolarized the HEK293 cell resting membrane potential by ∼-25 mV (Figs. 7A-C) (RMP = -76 ± 3 mV and -40 ± 7 mV, for TREK-1^CG*^ expressing cells and untransfected cells, respectively), consistent with substantial basal TREK-1^CG*^ activity. By contrast, TREK-1^CG*^ A286F expressing cells had basal currents at 0 mV that were similar to untransfected cells (90 ± 10 pA and 70 ± 20 pA, respectively) (Fig. 7A) and an unperturbed RMP (RMP = -45 ± 5 mV) (Figs. 7A-C). Applying CAT335 to TREK-1^CG*^ expressing cells resulted in a ∼8.5-fold current increase (5580 ± 1135 pA at 0 mV, Fig. 7C) but had a limited effect on RMP (ΔRMP = -5 mV, Fig. 7B) as the TREK-1^CG*^ basal activity had already hyperpolarized the cells. CAT335 application to TREK-1^CG*^ A286F expressing cells induced smaller leak currents (1556 ± 354 pA at 0 mV) but caused larger RMP changes (ΔRMP = -27 mV). Notably, CAT335 did not affect the leak currents and RMP of control cells (Fig. 7B). Hence, the data show that the CATKLAMP chemogenetic pairs, exemplified by CAT335 and TREK-1^CG*^ A286F act as a hyperpolarization switch that rapidly clamps the target cell near the potassium equilibrium potential.

**Figure 7.**
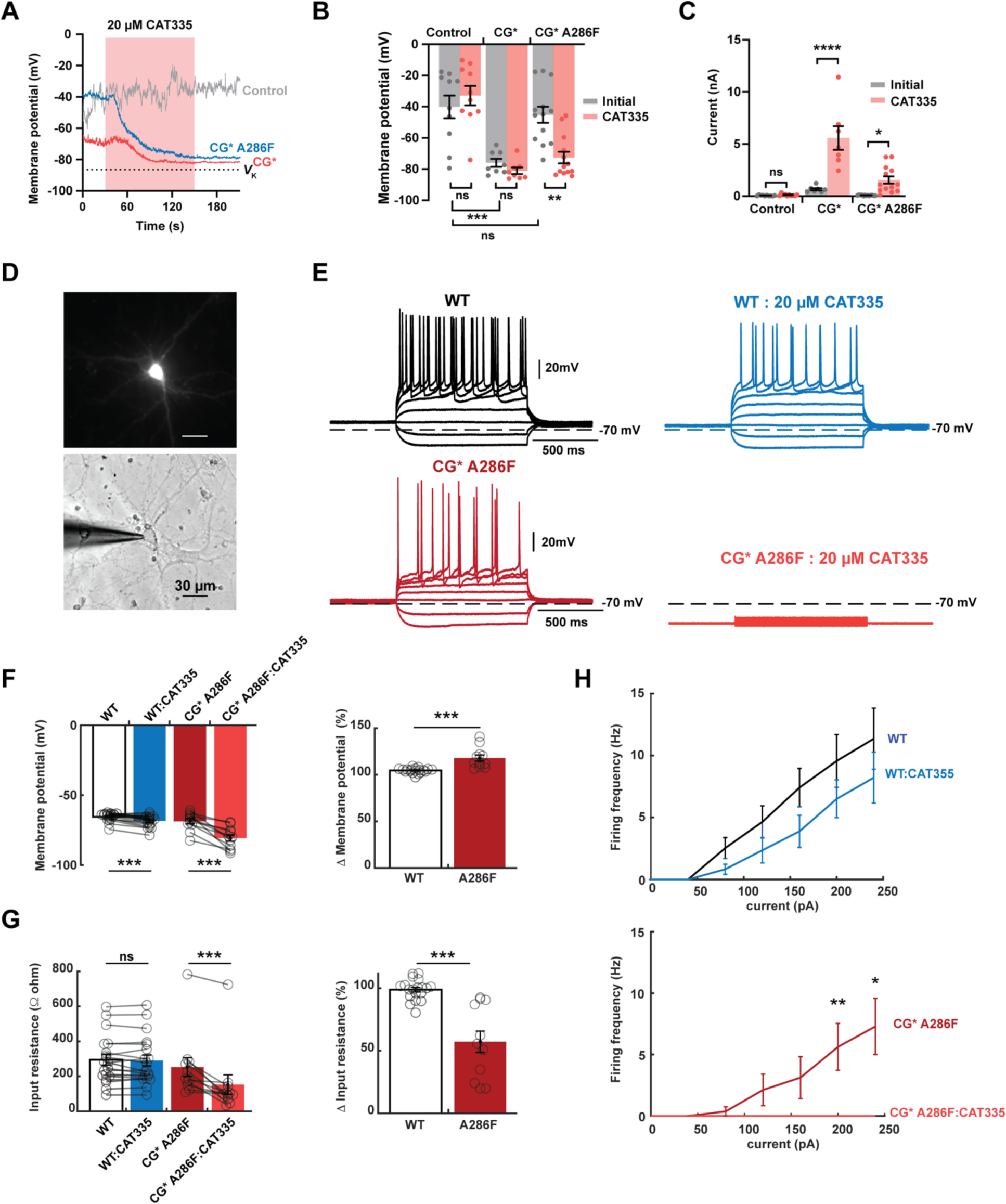
CATKLAMP activation with CAT335 hyperpolarizes cells. **A,** Exemplar gap-free, I=0 recordings of untransfected HEK293 (control, gray) or HEK293 cells expressing TREK-1^CG*^ (CG*) (red) or TREK-1^CG*^ A286F (CG* A286F) (blue). Red box indicates 20 µM CAT335 application. V_k_ indicates the K^+^ reversal potential. **B,** Change in Vm before (gray) and after (red) 20 µM CAT335 application (n= 8-13) for control, TREK-1^CG*^ (CG*), and TREK-1^CG*^ A286F (CG* A286F) cells. **C,** Whole-cell currents before (gray) and after 20 μM CAT335 application (red) (n=7-13) for cells from ‘B’. Significance was determined using either paired t-tests (for panel B) or unpaired t-tests (for panel C) where n.s. p>0.12; * p<0.033; **p<0.0021; *** p<0.0002; **** p<0.0001. **D,** Fluorescence (top) and bright field (bottom) image of exemplar mouse primary hippocampal neuron expressing TREK1^CG*^ A286F. **E,** Exemplar neuron responses to current injection with 1s steps from -80pA to 200pA for wild-type neurons and TREK1^CG*^ A286F expressing neurons before (black and red, respectively) and after 20 µM CAT335 application (blue and orange, respectively) **F,** RMP changes for the indicated neurons showing absolute (left) and normalized changes per neuron (right). **G,** Input resistance changes for the indicated neurons showing absolute values (left) and normalized changes per neuron (right). **H,** CAT335 effects on neuronal firing frequency in response to 1 s current steps for wild-type (top) and TREK1^CG*^ A286F neurons (bottom). For panels **F-H**, * p < 0.05, ** p<0.01, ***p<0.001 using a two-tailed Wilcoxon signed rank test for paired comparisons (**F** left panel, **G** left panel, and **H**) and Wilcoxon rank-sum for independent comparisons (**F** right panel, **G** right panel). Error bars represent S.E.M.

To test whether the CATKLAMP strategy could be applied to excitable cells, we developed a TREK-1^CG*^ A286F IRES GFP plasmid controlled by the human synapsin promoter and expressed this construct in primary mouse hippocampal neurons (Fig. 7D). Consistent with the HEK cell results, passive electric properties such as resting membrane potential and input resistance were not significantly different between wild-type neurons and TREK-1^CG*^ A286F expressing neurons (-68.7 ± 1.8 and -65.5 ± 0.7 mV; 294 ± 31 and 252 ± 54 MΩ respectively, Table S4). By contrast, application of 20 µM CAT335 to TREK-1^CG*^ A286F expressing cells caused dramatic changes in electrical activity (Fig. 7E). Perfusion of CAT335 onto TREK-1^CG*^ A286F expressing neurons substantially hyperpolarized the resting membrane potential (-68.7 ± 1.8 mV and -80.7 ± 2.2 mV, before and after CAT335, respectively) (Fig. 7F, Table S4), reduced input resistance (253 ± 54 MΩ and 153 ± 56 MΩ, before and after CAT335, respectively) (Fig. 7G, Table S4), and effectively suppressed action potential generation in response to current injection pulses (Fig. 7H). Notably, wild-type neurons were minimally affected by 20 µM CAT335 in all analyzed parameters, except for a mild hyperpolarization of ∼3mV (Figs. 7E-H, Table S4). Hence, these data demonstrate that the CATKLAMP strategy is a powerful means to silence neuronal firing.

### CATKLAMP strategy can be used in all TREK subfamily members

Previous studies showed that ML335 strongly activates K_2P_2.1(TREK-1) and K_2P_10.1(TREK-2), but not the third TREK subfamily member K_2P_4.1(TRAAK) unless this channel bears a Q258K mutation creating a cation-π interaction with the modulator upper ring^21^. Given the similarity between ML335 and CAT335 and similarities in TREK subfamily K_2P_ modulator pockets (Figs. 8A and B), we asked whether the CATKLAMP strategy could be extended to other TREK subfamily members. We made the K_2P_2.1(TREK-1) S131C equivalent change in K_2P_10.1(TREK-2) and K_2P_4.1(TRAAK), S152C and S91C, respectively, to create TREK-2^CG*^ and TRAAK^CG*^. Similar to K_2P_2.1(TREK-1) (Figs. 2A-C), K_2P_10.1(TREK-2) was insensitive to CAT335 whereas TREK-2^CG*^ responded to both ML335 and CAT335 (Figs. 8C and F, Table 1). Importantly, as with TREK-1^CG*^, 20 µM CAT335 activation of TREK-2^CG*^ was strong (*I*_CAT335_/*I*_0_ = 18 ± 2) and irreversible. By contrast, TRAAK^CG*^ did not respond to ML335 but could be irreversibly activated by 20 µM CAT335. The TRAAK^CG*^ response took longer to develop than that those observed for TREK-1^CG*^ or TREK-2^CG*^, showing some activation after 2 minutes (*I*_CAT335_/*I*_0_ = 4.2 ± 0.6) and reaching maximal activation after longer application (6 min) (*I*_CAT335_/*I*_0_ = 10.7 ± 1.0) (Figs. 8D and F, Table 1). Hence, the CATKLAMP approach is applicable to all members of the TREK subfamily.

**Figure 8:**
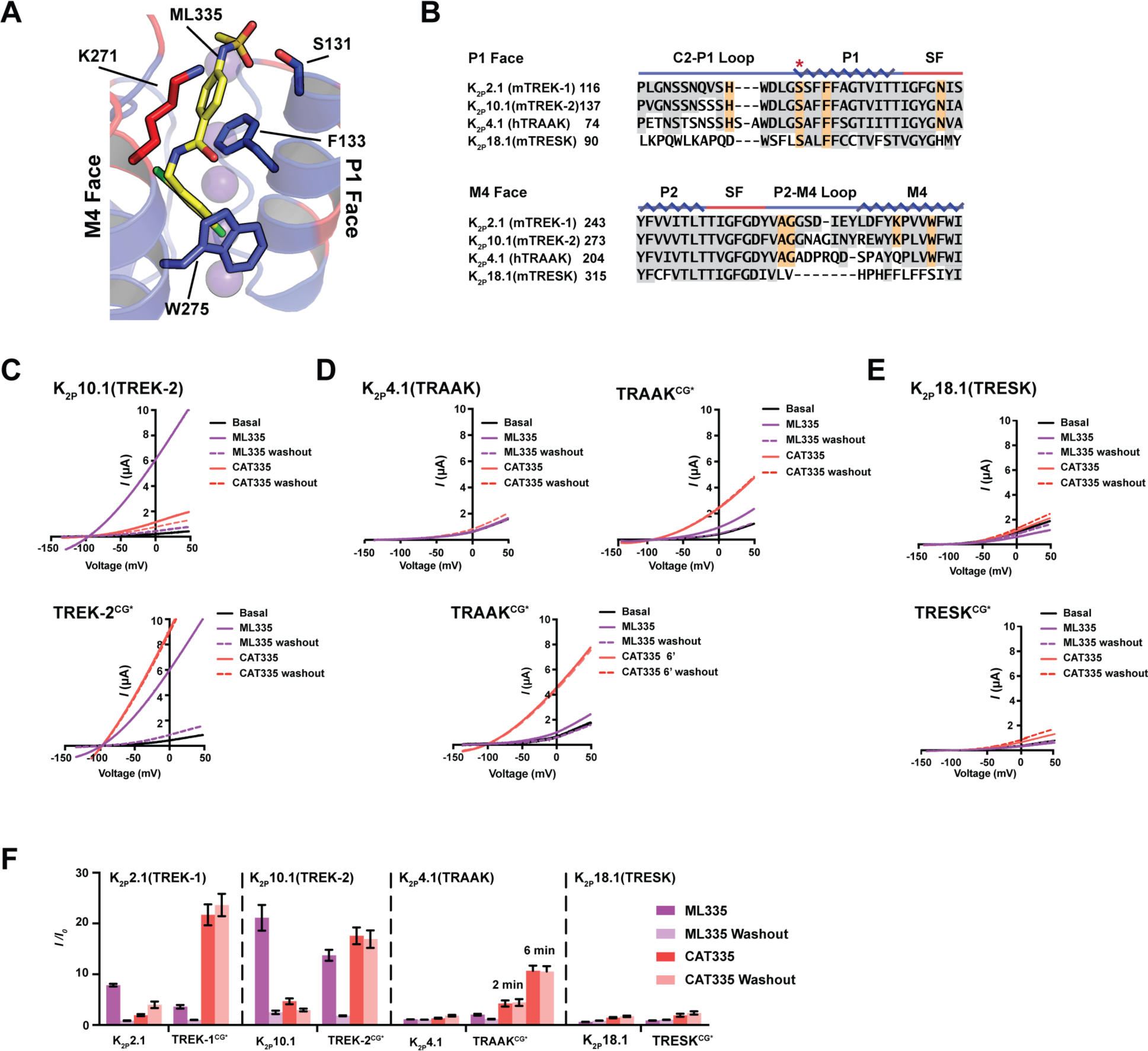
TREK subfamily members are compatible with the CATKLAMP strategy. **A,** View of the K_2P_ modulator pocket (PDB:6CQ8)^21^ showing TREK subfamily conservation. Blue (conserved), red (variable). M4 and P1 faces are labeled. Key residues for binding ML335 are shown as sticks. ML335 is yellow. **B,** Sequence comparison for mouse K_2P_2.1(TREK-1) (NP_034737.2), mouse K_2P_10.1(TREK-2) (NP_001303594.1), human K_2P_4.1(TRAAK) (NP_001304019.1), and mouse K_2P_18.1 (TRESK) (NP_997144.1). K_2P_ modulator pocket positions are orange. Conservation is grey. Red asterisk marks the chemogenetic SèC mutant site. Protein secondary structure is indicated. **C-E**, TEVC recordings showing responses to ML335 and CAT335 for **C,** K_2P_10.1(TREK-2) and TREK-2^CG*^, **D,** K_2P_4.1(TRAAK) and TRAAK^CG*^, and **E,** K_2P_18.1(TRESK) and TRESK^CG*^. **F,** Activation (*I/I_0_*) of indicated K_2P_s measured in *Xenopus* oocytes at 0 mV in response to 20 μM ML335 (magenta) and 20 μM CAT335 (red) (n=9-11). All responses in ‘C-F’ are at 2 minutes, unless indicated otherwise. Error bars show mean ± S.E.M.

A cation-π interaction between Lys271 on the K_2P_2.1 (TREK-1) M4 face of the K_2P_ modulator pocket and the K_2P_ activator upper ring (Fig. 8A) is crucial for TREK subfamily responses to ML335^21^. Because introducing this interaction in K_2P_4.1(TRAAK) via the Q258K mutation is sufficient to sensitize K_2P_4.1(TRAAK) to ML335^21^, we tested whether the same change would enhance TRAAK^CG*^activation by CAT335. Similar to K_2P_4.1(TRAAK) Q258K^21^, TRAAK^CG*^ Q258K was activated by 20 μM ML335 (*I*_ML335_/*I*_0_ = 3.7 ± 0.3) (Figs. S8A and C). This channel also showed a better response to a 2 minute application of CAT335 than its parent TRAAK^CG*^ (*I*_CAT335_/*I*_0_ = 7.4 ± 0.6 and 4.2 ± 0.6 after 2 min for TRAAK^CG*^ Q258K and TRAAK^CG*^, respectively) (Figs. 8D-F, S8A-C, Table 1), providing further support for the importance of the K_2P_ modulator pocket:agonist cation-π interaction. Examination of TREK-1^CG*^ channels bearing the converse change, K271Q, showed that like K_2P_2.1 (TREK-1) K271Q^21^, TREK-1^CG*^ K271Q was insensitive to ML335. However, TREK-1^CG*^ K271Q retained a strong CAT335 response (*I*_CAT335_/*I*_0_ = 8.4 ± 1.1) (Figs. S8A and C, Table 1). Together with the TRAAK^CG*^ results, these data provide clear evidence that the chemogenetic strategy can capture weak K_2P_ modulator pocket ligands.

To test the selectivity of this chemogenetic approach for enabling a K_2P_ to respond to CAT335, we implemented the same strategy on a K_2P_ outside of the TREK subfamily. We chose K_2P_18.1(TRESK), as this channel has numerous changes on the M4 face of the K_2P_ modulator pocket equivalent but maintains key P1 face positions (Ser131 and Phe133 equivalents) that could engage ML335 activators (Figs. 8A and B). TEVC experiments showed that neither K_2P_18.1(TRESK) or its chemogenetic partner having the S116C change, TRESK^CG*^, responded to ML335 (Figs. 8E and F) or to CAT335. Hence, the mere presence of an S131C equivalent is insufficient to render a K_2P_ responsive to CAT335. These observations together with the results from TRAAK^CG*^, demonstrate that both CAT335-competent modulator pocket and reactivity at the K_2P_2.1(TREK-1) Ser131 equivalent are necessary for producing a CAT335 responsive chemogenetic target channel and highlight the selectivity of CAT335 for the TREK subfamily.

## Discussion

Probing the chemical and electrical signals that power the brain, heart, and nervous system requires development of diverse means to control electrical signaling. Ongoing efforts to create optogenetic^32-37^ and chemogenetic^38,39^ approaches for modulating potassium channels highlight the potential to leverage control of this channel class to manipulate electrical excitability. As leak channels, K_2P_s are naturally poised to affect membrane potential and excitability^2,3,16^. Nevertheless, despite some effort to develop optogenetic tools using tethered light-switchable blockers^32^, the power of K_2P_s to manipulate membrane potential has been largely untapped^3,16^. Our CATKLAMP approach provides a new means to control K_2P_ function selectively, probe channel biophysical mechanisms of action, and a new strategy for clamping membrane potential and silencing electrical signaling.

The heart of K_2P_ function centers on the selectivity filter ‘C-type’ gate^16,40-42,47^. A range of physical and chemical gating cues influence interactions between the permeant potassium ions and selectivity filter to affect a ‘substrate-assisted’ transport mechanism in which ion occupancy and filter stability mutually affect ion permeation^16,21,23,40-43^. One control point for TREK subfamily K_2P_s is the K_2P_ modulator pocket, a cryptic binding site between the P1 and M4 helices where small molecule activators ML335 and ML402 act as interdomain wedges that stabilize the C-type gate active conformation and enable ion permeation^16,21,23^. CATKLAMP capitalizes on this mechanism by providing a covalent ligand for the K_2P_ modulator pocket, CAT335, that spares the wild-type channel but binds in an irreversible manner when the target subunit has a cysteine substitution at the K_2P_2.1(TREK-1) Ser131 position or its equivalent yielding irreversible channel activation (Figs. 2A-H, 8C-D and F).

Each TREK subfamily channel has two K_2P_ modulator pockets^16,21^. It has been unclear whether occupation of both is necessary to activate the channel or how a channel would respond to occupation of only one site. By exploiting the selectivity of the CATKLAMP ligand CAT335 for the modified target subunit, TREK^CG*^, we were able to address this question using concatenated versions of K_2P_2.1(TREK-1) in which one or both K_2P_ modulator pockets within the K_2P_ channel dimer were CAT335 responsive. The data show that occupation of a single K_2P_ modulator pocket by CAT335 is sufficient to evoke up to ∼80% of the maximal response of the channel to K_2P_ modulator pocket agonists (Fig. 4C). This activation is accompanied by a dramatic increase in channel open probability (Fig. 5P), similar to that seen for the non-covalent agonist ML335^23^. Hence, activation of a single subunit in the K_2P_ dimer is sufficient to activate the C-type gate. The observation that activation of a single subunit yields >50% of the expected maximal response for occupation of both binding sites underscores the cooperativity between the K_2P_ modulator pocket and selectivity filter and reinforces the idea that the K_2P_2.1(TREK-1) selectivity filter active conformation is influenced by both K_2P_ modulator pocket occupation and permeant ions^23^. Heterodimer formation is emerging as an important issue in K_2P_ channel functional diversification^12,52-55^. Notably, TREK subfamily members appear quite adept at making both intrafamily, ex. K_2P_2.1 (TREK-1)/K_2P_4.1 (TRAAK)^53,54^, and interfamily, ex. K_2P_2.1 (TREK-1)/K_2P_1.1 (TWIK-1)^56^, heterodimers. Our finding that occupation of one K_2P_ modulator pocket is sufficient to stimulate channel activity suggests that K_2P_ modulator pocket activators should be effective at activating K_2P_ heterodimers in which only one subunit is agonist sensitive. In this regard, demonstration that other TREK subfamily channels can be rendered CAT335 responsive (Fig. 8), opens a new tactic for using the CATKLAMP approach to probe the growing list of heterodimeric K_2P_s^55^.

The polysite nature of K_2P_ pharmacology raises the possibility for synergistic action among modulators acting on the various effector binding sites^16,57^. However, apart from the observed independence of the K_2P_ modulator pocket (MP) and Keystone inhibitor site located under the K_2P_ cap^20^, the degree of coupling among the various modulator binding sites has been largely unexplored. BL-1249 activates the K_2P_ C-type gate^19,48^ by binding at the fenestration site (FS) located under the selectivity filter^16,19^. Even though the M4 helix that forms part of the FS is a well-established effector C-type gate function^19,24,40,47,48,50,51^, whether the FS and K_2P_ modulator pocket site are coupled has been unclear. Our studies revealed synergistic activation when ML335 and BL-1249 were applied simultaneously (Fig. S7 A and C, Table S3) and an unexpected priming behavior in which application of the FS agonist BL-1249 enhanced subsequent MP site CAT335 responses (Fig.6 E-J) providing clear evidence of coupling and synergy between the two sites. These observations also corroborate the idea that M4 helix movement controls fenestration site access^19,24^ and impacts the conformation of the selectivity filter and surrounding structure^21,23,40,47,48^. They further are in line with the fact that the N-terminal end of the M4 helix forms part of the K_2P_ modulator pocket^21,23^. This synergy between the FS and MP sites opens the possibility for developing means to affect K_2P_ function through simultaneous targeting of the MP and FS sites and requires further investigation.

The ability of covalent small molecule adducts to control protein function is well established for drugs^58^ and a variety of chemical tools designed to probe diverse biological targets^58-61^. Covalent drugs and chemical probes exploit the combination of binding affinity for their targets and chemical reactivity to achieve selectivity^60^. Here, we used the same principles to create a covalent activator selective for engineered versions of TREK subfamily members. Because potassium channel activation quiets electrical activity^31^, potassium channel activation has an inhibitory effect. The use of the CATKLAMP system in both HEK293 and transfected neurons establishes its utility as a means for providing a rapid and stable clamp of the cell membrane potential at the potassium reversal potential (Fig. 7A and F) to silence neuronal firing (Fig. 7E). As an irreversible, covalent agonist that works in the extracellular oxidative environment, CATKLAMP effects would be expected to persist for the life of the channel on the plasma membrane. The consequent long lived inhibition of excitation, similar to the effects described for the light-activated potassium channel BLINK2^37^, could be advantageous for studies where persistent inactivation of a particular class of neurons is of interest, such as in pain or behavior studies.

The combination of binding affinity and covalent modification allows one to probe target protein sites that might otherwise have low ligand binding affinity. This principle is exemplified in cysteine-disulfide tethering for ligand discovery^62-64^ and is evident in the differential responses to CAT335 within the TREK subfamily. Even though K_2P_4.1(TRAAK) is not sensitive to ML335 due to the lack of a key K_2P_ modulator pocket cation-π interaction^21^, its chemogenetic version, TRAAK^CG*^, can be irreversibly activated by CAT335. In line with its M335-insenstivity, the TRAAK^CG*^ response to CAT335 is slower than the CAT335 responses of the ML335-sensitive channels TREK-1^CG^* and TREK-2^CG*^ (Fig. 8F). Further, TRAAK^CG*^ reactivity to CAT335 can be enhanced by supplying the cation-π interaction (Figs. 8D-F, S8A-C, Table 1) providing additional evidence that the K_2P_4.1(TRAAK) K_2P_ modulator pocket is accessible to small molecule activators^21^. Importantly, the mere presence of a reactive cysteine in this part of the channel is not sufficient, as making the analogous change in the distantly related K_2P_18.1 (TRESK) channel failed to confer CAT335 sensitivity. Together, these results underscore that covalent engagement is able to capture weak binding events driven by the ML335 core and suggest that the chemogenetic approach may be a fruitful way to broaden the pharmacology of the K_2P_ modulator pocket and search for new ligands among different K_2P_ subtypes. Our finding that the K_2P_2.1 (TREK-1) K_2P_ modulator pocket residue Ser131 could be modified by the ML335 acrylamide derivative ML336 (Fig. 1B) was a surprise given the generally low reactivity of hydroxyls at neutral pH. This observation, together with the success of using a cysteine mutation at this position as a target for maleimides such as CAT335 (Figs. 3A and 4-8), and the apparent slow, but natural reactivity of the native Ser131 suggests that it may be possible to develop CAT335 derivatives that could target the native channel. Additionally, as modification of intracellular cysteines by electrophilic compounds in mustard oil is a natural process for activation of the wasabi receptor ion channel TRPA1^65-68^, there may be some natural, modifier of Ser131 that exploits its reactivity and affects K_2P_ function.

Chemogenetic tools offer precise chemical control of protein function that is orthogonal to native protein-ligand interactions and are well developed for G-protein coupled receptors and some ligand-gated ion channels^25,26^. Our development of the CATKLAMP covalent modification system provides a generalizable concept for the ion channel chemical biology toolbox^69^ that should be applicable to any ion channel whose action is controlled by a small molecule ligand. We note that CATKLAMP may offer a means to label and track TREK subfamily channels by modifying the CAT335 scaffold with a second chemical handle using strategies similar to those used for saxitoxin-maleimide derivatives and voltage-gated sodium channels^70,71^. We also anticipate that leveraging the properties of this system to capture ligands, such as with tethering approaches^62-64^, should enable identification of new molecules to control K_2P_ function. Moreover, because K_2P_ channel activation is well suited to silence neuronal activity and such silencing should be a powerful way to probe networks in the brain^44^, we expect that implementation of the CATKLAMP strategy will facilitate new means to probe processes such a pain responses and behavior.

## Supporting information

Supplementary Figs. S1-S8, Scheme 1, Tables S1-S4, and references

## Acknowledgements

We thank F. Abderemane-Ali and S. Jang for electrophysiology guidance, and K. Brejc and D. Zuniga for comments on the manuscript. This work was supported by a UCSF Weill Trailblazer Award in Neurosciences to A.R.R. and D.L.M., NIH grants T32HL007731 to P.E.D., R01NS119826 and R01NS107506 to E.Y.I., R01MH116278 to A.R.R. and D.L.M., and R01-MH093603 to D.L.M..

## Author Contributions

P.E.D., M.L., A.R.R., and D.L.M. conceived the study and designed the experiments. M.L. H.L., and A.M. expressed, purified, and crystallized the proteins, collected diffraction data, and determined the structures. P.E.D., X.E.-H., and C.B. synthesized and purified the compounds. P.E.D. and M.L. performed functional studies. P.R.F.M. and H.B. designed neuronal expression vectors, and carried out and analyzed neuron recordings. P.E.D., H.L., A.M., M.L., P.R.F.M., H.B., E.Y.I., A.R.R. and D.L.M. analyzed the data. E.I.Y., A.R.R. and D.L.M. provided guidance and support. P.E.D., H.L., A.M., M.L., P.R.F.M., H.B., E.Y.I., A.R.R. and D.L.M wrote the paper.

## Materials and correspondence

Correspondence should be directed to A.R.R. or D.L.M. Requests for materials should be directed to D.L.M.

## Data and materials availability

Coordinates and structures factors are deposited in the RCSB and will be released immediately upon publication.

## Materials and Methods

### Molecular Biology

Mouse K_2P_2.1(TREK-1) (Genbank accession number: NP_034737.2), mouse K_2P_10.1(TREK-2) (NM_001316665.1), mouse K_2P_4.1(TRAAK) (NM_008431.3), human K_2P_4.1(TRAAK) (NM_033310.2), and mouse K_2P_18.1(TRESK) (NM_207261.3), and an engineered mouse K_2P_2.1 (TREK-1), denoted K_2P_2.1 ^21^, encompassing residues 22-322 and bearing the following mutations: K84R, Q85E, I88L, A89R, Q90A, A92P, N95S, S96D, T97Q, N119A, S300A, and E306A were used for this study. Point mutations were introduced by site-directed mutagenesis using custom primers and confirmed by sequencing. For studies using *Xenopus* oocytes, K_2P_ channels were subcloned into a previously reported pGEMHE/pMO vector^40^. For expression in HEK293 cells, mouse K_2P_2.1(TREK-1) (NP_034737.2) and mutants were expressed from a previously described pIRES2-EGFP vector in HEK293 cells^40,72^. K_2P_2.1(TREK-1) tandems were constructed by connecting the open reading frames for the individual subunits with a linker encoding the AAAGSGGSGGSGGSSGSSGS sequence. Tandems used for HEK293 cells included an N-terminal HA tag (YPYDVPDYA) on the first subunit.

For expression in neurons, the CG*A286F 5-IRES-GFP DNA fragment was extracted from the pIRES2-eGFP_A2865F-S131C-mTREK-1 plasmid and inserted into a custom pCDNA3.1(+) vector containing the human synapsin promoter, hSyn, in place of the standard pCDNA CMV promoter and in which the NeoR/KanR cassette was removed and the AmpR cassette was used for bacterial selection creating a plasmid named hSyn-CG*A286F-IRES-GFP. The DNA insertion and vector were amplified using Phusion Hot Start II DNA Polymerase (Thermo Scientific Cat #F549L) and custom forward and reverse primers from Integrated DNA Technologies. Primers for insert amplification contained homology to the target vector in order to generate amplified sequence ends compatible with downstream Gibson Assembly. TREKA2865-IRES-GFP was inserted into the vector using in-house Gibson enzyme master mix for 2-piece Gibson Assembly^73^. Primers used: reverse vector: ggtaccagatctgaattcgactgcg; forward vector: aggatcccgggtggcatccctgtgac; Reverse insert: cgcagtcgaattcagatctggtaccatgtacccatacgacgtgccagac; forward insert: cacagggatgccacccgggatcctttacttgtacagctcgtccatgccgag.

### Protein expression

K_2P_2.1_cryst_ and K_2P_2.1_cryst_ S131C bearing a C-terminal green fluorescent protein (GFP) and His_10_ tag were expressed from a previously described *Pichia pastoris* pPICZ vector^21^. Plasmids were linearized with *PmeI* and transformed into *P. pastoris* SMD1163H by electroporation. Multi-integration recombinants were selected by plating transformants onto yeast extract peptone dextrose sorbitol (YPDS) plates having increasing concentrations of zeocin (0.5–2 mg ml^−1^). Expression levels of individual transformants were evaluated by FSEC as previously described^74^. Large-scale expression was carried out in a 7L Bioreactor (Labfors5, Infors HT). First, a 250 ml starting culture was grown in buffered minimal medium (2× YNB, 1% glycerol, 0.4 mg L^−1^ biotin, 100 mM potassium phosphate, pH 6.0) in shaker flasks for two days at 29 °C. Cells were pelleted by centrifugation (3,000*g*, 10 minutes, 20 °C) and used to inoculate the bioreactor. Cells were grown in minimal medium (4% glycerol, 0.93 g L^−1^ CaSO_4_·2H_2_O, 18.2 g L^−1^ K_2_SO_4_, 14.9 g L^−1^ MgSO_4_.7H_2_O, 9 g l^−1^ (NH_4_)_2_SO_4_, 25 g L^−1^ Na^+^ hexametaphosphate, 4.25 ml L^−1^ PTM_1_ trace metals stock solution prepared accordingly to standard Invitrogen protocol) until the glycerol in the fermenter was completely metabolized marked by a spike in pO_2_ (around 24 h). Fed-batch phase was then initiated by adding a solution of 50% glycerol and 12 ml L^−1^ of trace metals at 15-30% of full pump speed until the wet cell mass reached approximately 250 g L^−1^ (around 24 h). pO_2_ was measured continuously and kept at a minimum of 30%. Feed rate was automatically regulated accordingly. pH was maintained at 5.0 by the addition of a 30% ammonium hydroxide solution.

After the fed-batch phase was completed, cells were then starved to deplete glycerol by stopping the feeder pump until a pO_2_ spike appeared. After starvation, the temperature was set to 27 °C, and the induction was initiated with addition of methanol in three steps: (1) initially, the methanol concentration was kept at 0.1% for 2 h in order to adapt the cells; (2) methanol concentration was then increased to 0.3% for 3 h; and (3) methanol was then increased to 0.5% and expression continued for 48–60 h. Cells were then pelleted by centrifugation (6,000*g*, 1 h, 4 °C), snap frozen in liquid nitrogen, and stored at −80 °C.

### Protein purification

In a typical preparation, 30-50 g of cells were broken by cryo-milling (Retsch model MM400) in liquid nitrogen (3 × 3 min, 25 Hz). All subsequent purification was carried out at 4 °C. Cell powder was added at a ratio of 1 g cell powder to 3 ml lysis buffer (200 mM KCl, 21 mM OGNG (octyl glucose neopentyl glycol, Anatrace), 30 mM HTG (*n*-heptyl-β-D-thioglucopyranoside, Anatrace), 0.1% CHS, 0.1 mg ml^−1^ DNase 1 mM PMSF, 100 mM Tris-Cl, pH 8.2). Membranes were extracted for 3 h with gentle stirring followed by centrifugation (100,000*g*, 45 minutes at 4 °C).

Solubilized proteins were purified by affinity chromatography using batch purification. Anti-GFP nanobodies were conjugated with CNBr Sepharose beads (GE Healthcare, #17-0430-02). The resin was added to the cleared supernatant at a ratio of 1 ml of resin per 10 g of cell powder and incubated at 4 °C for 3 h with gentle shaking. Resin was collected into a column and washed with 10 column volumes (CV) of buffer A (200 mM KCl, 10 mM OGNG, 15 mM HTG, 0.018% CHS, 50 mM Tris-Cl, pH 8.0) followed by a second wash step using 10 CV of buffer B (200 mM KCl, 5 mM OGNG, 15 mM HTG, 0.018% CHS, 50 mM Tris-Cl, pH 8.0). The resin was then washed with additional 10 CV of buffer C (200 mM KCl, 3.85 mM OGNG, 15 mM HTG, 0.0156% CHS, 50 mM Tris-Cl, pH 8.0). On column cleavage of the affinity tag was achieved by incubating the resin with buffer C supplemented to contain 350 mM KCl, 1 mM EDTA, and 3C protease at ratio of 50:1 resin volume:protease volume.^46^ The resin was incubated overnight at 4 °C. Cleaved sample was collected and the resin washed with 2 CV of SEC buffer (200 mM KCl, 2.1 mM OGNG, 15 mM HTG, 0.012% CHS, 20 mM Tris-Cl, pH 8.0). Purified sample was concentrated and applied to a Superdex 200 column pre-equilibrated with SEC buffer. Peak fractions were evaluated by SDS-PAGE (15% acrylamide) for purity, pooled and concentrated.

### Crystallization and refinement

For K_2P_2.1_cryst_ ML336 crystals, ML336 was dissolved in 100% DMSO at a concentration of 500 mM, diluted 1:100 in SEC buffer to 5 mM concentration, and then mixed to dissolve the compound completely. 12 mg ml^−1^ K_2P_2.1_cryst_ then mixed 1:1 with the ML336-SEC buffer solution to achieve a final concentration of 6 mg ml^−1^ and 2.5mM of K_2P_2.1_cryst_ and ML336, respectively. This mixture was then incubated for 2h on ice prior to crystallization using hanging-drop vapor diffusion at 4 °C and mixture of 0.2 μl of protein and 0.1 μl of precipitant over 100 μl of reservoir containing 20–25% PEG400, 200 mM KCl, 100 mM HEPES pH 8.0, 1 mM CdCl_2_. Crystals appeared in 12 h and grew to full size (200–300 μM) in about a week. Crystals were cryoprotected with buffer D (200 mM KCl, 0.2% OGNG, 15 mM HTG, 0.02% CHS, 100 mM HEPES pH 8.0,1 mM CdCl_2_) with 5% step increase of PEG400 up to a final concentration of 38% and flash-frozen in liquid nitrogen.

TREK-1^CG^*:CAT335 crystals, TREK-1^CG*^:CAT335a crystals, and TREK-1^CG^*:ML335 crystals grew in the similar conditions as the TREK-1:ML336 complex, but CAT335 and CAT335a were dissolved in 100% DMSO at a concentration of 20 mM and ML335 was dissolved in 100% DMSO at a concentration of 500 mM. Each ligand was diluted in SEC buffer and mixed 1:1 volume ratio to TREK-1^CG*^ previously concentrated to 10 mg ml^−1^ to achieve final ligand:protein molar ratios of 1.1:1 CAT335, 2:1 CAT335a, and 5:1 ML335. These mixtures were incubated for 2h on ice prior to crystallization using hanging-drop diffusion at 4 °C and a mixture of 0.2 μl of protein and 0.1 μl of precipitant over 100 μl of reservoir containing 20–25% PEG400, 200 mM KCl, 100 mM HEPES pH 7.0 or 7.5, and 1 mM CdCl_2_. Crystals appeared in 12 h and grew to full size (200-300 μM) in about a week. Crystals were cryoprotected with buffer D (200 mM KCl, 0.2% OGNG, 15 mM HTG, 0.02% CHS, 100 mM HEPES pH 7.0 or 7.5, and 1 mM CdCl_2_) with 5% step increase of PEG400 up to a final concentration of 38% and flash-frozen in liquid nitrogen.

Datasets for K_2P_2.1–ligand complexes were collected at 100 K using synchrotron radiation at APS GM/CAT beamline 23-IDB/B Chicago, Illinois using a wavelength of 0.9779 Å, processed with XDS^75^, scaled and merged with Aimless^76^. Final resolution cut-off was 2.9 Å (K_2P_2.1_cryst_:ML336) and 3.0 Å (K_2P_2.1^CG*^:CAT335, K_2P_2.1^CG*^:CAT335a, and K_2P_2.1^CG*^:ML335), using the CC_1/2_ criterion^77,78^. Structures were solved by molecular replacement using the K_2P_2.1_cryst_ structure (PDB: 6CQ6) as search model. Several cycles of manual rebuilding, using COOT^79^ and refinement using REFMAC5^80^ and PHENIX^81^ were carried out to improve the electron density map. Ramachandran restraints were employed during refinement. Model quality was assessed using Molprobity^82^.

### Patch-clamp electrophysiology

50% confluent cells were co-transfected (in 35-mm diameter wells) with 10-100 ng of the K_2P_2.1 plasmid and 400 ng of an eGFP plasmid (for visualization) using LipofectAMINE 2000 (Invitrogen) for 24 h, after which the cells were either plated onto coverslips or the media exchanged with fresh media. Cells were plated onto Matrigel coated coverslips (BD Biosciences) 1-2 hours before experiments.

Whole-cell patch-clamp experiments with K_2P_2.1(TREK-1) and mutants were carried out 24-48 h after transfection. Acquisition and analysis for voltage-clamp experiments were performed using pCLAMP10 and an Axopatch 200B amplifier (Molecular Devices). Current-clamp experiments were acquired using an Axopatch 700B amplifier (Molecular Devices). Pipettes were pulled using a Flaming/Brown micropipette puller (P-97, Sutter Instruments) and flame-polished using a MF-830 microforge (Narishige). Electrode resistance after filling with pipette internal solution ranged from 2 to 5 MΩ. Currents were low-pass filtered at 2 kHz and sampled at 10 kHz. Pipette solution contained the following: 145 mM KCl, 3 mM MgCl_2_, 5 mM EGTA and 20 mM HEPES (pH 7.2 with KOH). Bath solution contained the following: 145 mM NaCl, 5 mM KCl, 1 mM CaCl_2_, 3 mM MgCl_2_ and 20 mM HEPES (pH 7.4 with NaOH). K_2P_2.1 currents were elicited by a 1 s ramp from –100 to +50 mV from a –80 mV holding potential. Current-clamp recordings were taken with no holding current (I=0). For K_2P_2.1(TREK-1)CG*, whole-cell patches with basal currents ≤2000 nA at 0 mV were used. ML335 or CAT335 were applied at 20 μM in bath solution and were perfused at 3 ml min^-1^ until potentiation stabilized (1-3 min). Data were analyzed using Clampfit 11 and GraphPad Prism 9.

Single-channel recordings of K_2P_2.1(TREK-1) tandem mutants were recorded in cell-attached mode using HEK293 cells 48 h after transfection. Acquisition and analysis was preformed using pCLAMP9 and an Axopatch 200B amplifier. Pipettes were pulled using a laser-based micropipette puller (P-2000, Sutter Instruments). Electrode resistance after filling with pipette internal solution ranged from 6 to 10 MΩ. Pipette solution contained the following: 150 mM KCl, 3.6 mM CaCl_2_, 10 mM HEPES (pH 7.4 with KOH). The bath solution was identical to the pipette solution. P_o_ was measured from 30-120 s recordings of K_2P_2.1(TREK-1) activity holding at -100 mV. Channel conductance of K_2P_2.1(TREK-1) tandems were measured from 30-60s recordings at either -100 mV or +50 mV holding potential. Recordings of CAT335 activated channels were obtained by bath applying 20 μM CAT335 for 2 min to the coverslip before transferring to the recording chamber. This was necessary as application of CAT335 via perfusion or through the pipette solution activated slowly (>5 min), making it difficult to measure P_o_ from fully activated channels. BL-1249 was applied at 20 μM in the bath solution via perfusion at 3 mL min^-1^ (2-4 min). Data were analyzed using Clampfit 11 and GraphPad Prism 9.

### Two-electrode voltage-clamp (TEVC) electrophysiology

mRNA for oocyte injections was prepared from linearized plasmid DNA (linearized with AflII) using mMessage Machine T7 Transcription Kit (Thermo Fisher Scientific). RNA was purified using RNEasy kit (Qiagen) and stored as stocks and dilutions in RNAse-free water at -80 °C.

*Xenopus laevis* oocytes were harvested according to UCSF IACUC Protocol AN129690 and digested using collagenase (Worthington Biochemical Corporation, #LS004183, 0.8-1.0 mg/mL) in Ca^2+^-free ND96 (96 mM NaCl, 2 mM KCl, 3.8 mM MgCl_2_, 5 mM HEPES pH 7.4) immediately post-harvest, as previously reported.^31,33^ Oocytes were maintained at 18 °C in ND96 (96 mM NaCl, 2 mM KCl, 1.8 mM CaCl_2_, 2 mM MgCl_2_, 5 mM HEPES pH 7.4) supplemented with antibiotics (100 units ml^-1^ penicillin, 100 µg ml^-1^ streptomycin, 50 µg ml^-1^ gentimycin) and used for experiments within one week of harvest. Defolliculated stage V-VI oocytes were microinjected with 0.5-36 ng mRNA and currents were recorded within 16-48 h hours of injection.

For recordings, oocytes were impaled with borosilicate recording microelectrodes (0.4–2.0 MΩ resistance) backfilled with 3 M KCl and were subjected to constant perfusion of ND96 at a rate of 3 ml min^−1^. Recording solutions containing K_2P_ activators were prepared immediately prior to use from DMSO stocks (20-100 mM), with final DMSO concentrations of 0.1%. Currents were evoked from a -80 mV holding potential followed by a 300 ms ramp from -150 mV to +50 mV. Data were recorded using a GeneClamp 500B amplifier (MDS Analytical Technologies) controlled by pCLAMP8 software (Molecular Devices), and digitized at 1 kHz using Digidata 1332A digitizer (MDS Analytical Technologies). For each recording, control solution (ND96) was perfused over a single oocyte until current was stable before switching to solutions containing the test compounds at various concentrations. Representative traces and dose response plots were generated in GraphPad Prism 9 (GraphPad Software, Boston) .

### Primary neuronal culture

Primary hippocampal neuronal cultures were prepared from P0-P1 mouse brains according to UC Berkeley IACUC protocol AUP-2015-06-7720-2. Briefly, neonate hippocampi were dissected in ice cold saline solution (HBSS, HEPES 10mM, Sodium Pyruvate 1mM, D-glucose 10 mM, pH 7.4 with NaOH) and digested with TrypsinLE (Gibco) for 15 minutes at 37° C. Following digestion, tissue was washed 3 times with saline solution and gently dissociated with a P1000 pipette tip (6-8 strokes). Dissociated tissue was strained with a 40 µm strainer (Falcon). Neurons were then plated onto 18mm glass coverslips coated with 1 mg ml^-1^ Poly-L-Lysine (Dissolved in borate buffer, pH 8.5) and maintained in culture media composed of Neurobasal-A medium (Gibco/Thermo Fisher), 0.1% B27 supplement (Gibco/Thermo Fisher), 1.2% Glutamax (Gibco/Thermo Fisher), and 1% PenStrep (Thermo Fisher). Each pair of hippocampi cells were plated onto ∼3 coverslips, yielding approximately 100,000 neurons per coverslip.

### Neuronal transfection

At 6 days *in vitro* (DIV), hippocampal cultures were transfected with hSyn-CG*A286F-IRES-GFP using Lipofectamine transfection (Invitroen). Briefly, Neurobasal-A was equilibrated at 37°/5% CO_2_ prior to transfection procedure. 500 ng of plasmid DNA and 1.5 µl Lipofectamine2000 (Invitrogen) were separately incubated in 30 µl Neurobasal-A (Gibco/Thermo Fisher) for 5 minutes at room temperature. The two solutions were then combined and further incubated for 20 minutes. Afterward, the media on neuronal culture was removed and saved, and 500 µl fresh Neurobasal-A was added to each well. The combined Lipofectamine/DNA mixture was then added dropwise onto each well. Neurons were incubated at 37°C for 2 hours. Transfection solution was aspirated and replaced with a solution at 37°C comprising ⅔ saved conditioned media and ⅓ fresh culture media.

### Neuronal electrophysiology and data analysis

Whole-cell current clamp recordings were carried out with an Axopatch 200B amplifier and CV203BU head stage (Molecular Devices). WinWCP software (developed by John Dempster, University of Strathclyde) was used to control Digidata 1440A acquisition board (Molecular Devices), with sampling rate at 20KHz, Bessel filtered at 4KHz. Neuronal cultures were used for electrophysiological recordings at 14-21 DIV. Coverslips were transferred to the recording chamber (total volume of ∼2ml) and perfused with extracellular solution at room temperature composed of (in mM): 139 NaCl, 2.5 KCl, 10 HEPES, 10 D-glucose, 2 CaCl_2_, 1.3 MgCl_2_ (pH 7.3 balanced with NaOH). Extracellular solution was supplemented with the synaptic blockers 5-phosphono-D-norvaline (AP5) (50 µM, Cayman), Gabazine (20 µM, Cayman), and 2,3-Dioxo-6-nitro-1,2,3,4-tetrahydrobenzo[f]quinoxaline-7-sulfonamide (NBQX) (10 µM, Cayman) to reduce spontaneous network activity. Borosilicate glass pipettes with resistance of 4-7 MΩ were filled with intracellular solution composed of (in mM): 105 K^+^ Gluconate, 30 KCl, 10 HEPES, 10 Phosphocreatine-Na_2_, 4 ATP-Mg, 0.3 GTP-NaH_2_0, 1 EGTA (pH=7.3, balanced with KOH, 284mOsm). Liquid junction potential was not corrected.

Pyramidal shaped neurons were selected for recordings, and GFP fluorescence was used to confirm CG*A286F expression. Wild-type neurons were recorded from either non transfected coverslips, or from transfected coverslips in which neurons did not express any GFP fluorescence. After establishing whole-cell configuration, passive and firing properties were measured in current-clamp mode as done previously^83^. Baseline excitability was measured with a family of 1s current steps starting at -80pA with 40pA increments. Afterward, the effect of CAT335 on resting membrane potential (Vm) and input resistance (Rin) were measured using a current clamp recording protocol composed of a 1s 0pA, 1s -60pA hyperpolarizing step, and a 1s depolarizing step. CAT335 (20 µM) was perfused for ∼5 minutes. The effects on Vm and Rin for most CG*A286F neurons were normally observable within 2 minutes of perfusion. Afterward, the same family of current steps was applied to verify effects on excitability.

Data were analyzed using custom written MATLAB 2022a scripts (Mathworks, Natick, MA). Action potentials were detected using the built-in function findpeaks. Data normality was tested using the Kolmogorov Smirnov test, and non-parametric statistical tests, like the Mann-Whitney U test, were preferentially applied. Statistical values are shown as mean ± standard error of the mean (SEM).

### Chemical Synthesis - Materials and Instrumentation

Chemical reagents and solvents (dry) were purchased from commercial suppliers and used without further purification. Synthesis of ML335 was carried out as previously reported.^31^ Thin layer chromatography (TLC) (Silicycle, F254, 250 μm) was performed on glass backed plates pre-coated with silica gel and was visualized by fluorescence quenching under UV light. Column chromatography was performed on Silicycle Sili-prep cartridges using a Biotage Isolera Four automated flash chromatography system. NMR spectra were measured using a Varian INOVA 400 MHz spectrometer (with 5 mm QuadNuclear Z-Grad probe). Chemical shifts are expressed in parts per million (ppm) and are referenced to CDCl_3_ (7.26 ppm, 77.0 ppm) or DMSO (2.50 ppm, 40 ppm). Coupling constants are reported as Hertz (Hz). Splitting patterns are indicated as follows: s, singlet; d, doublet; t, triplet; q, quartet, dd, doublet of doublet; m, multiplet. LC-MS was carried out using a Waters Micromass ZQTM, equipped with Waters 2795 Separation Module and Waters 2996 Photodiode Array Detector and an XTerra MS C18, 5 μm, 4.6 x 50 mm column at ambient temperature. The mobile phases were MQ-H_2_O with 0.1% formic acid (eluent A) and HPLC grade methanol with 0.1% formic acid (eluent B). Signals were monitored at 254 over 15 min with a gradient of 10-100% eluent B.

### Chemical Syntheses

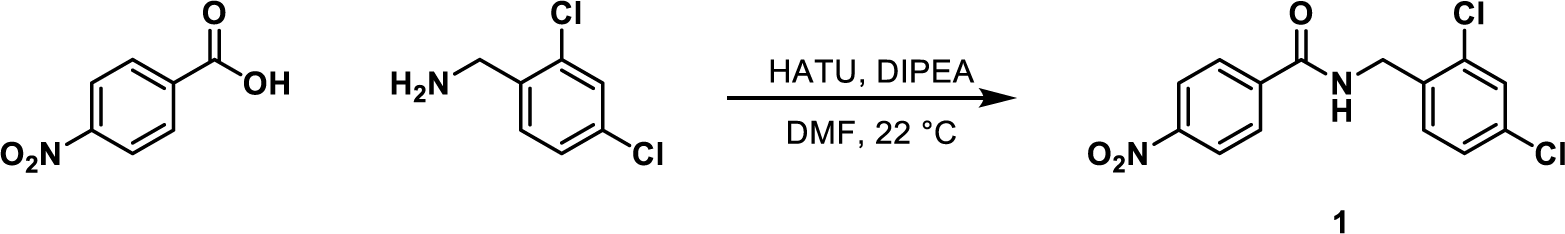

### Synthesis of N-[(2,4-dichlorophenyl)methyl]-4-nitrobenzamide, 1

A round-bottom flask was charged with 4-nitrobenzoic acid (2.00 g, 12.0 mmol), 2,4-dichlorobenzylamine (1.77 mL, 13.2 mmol), and HATU (5.00 g, 13.2 mmol). Anhydrous DMF (20 mL) and anhydrous diisopropylethylamine (4.17 mL, 24.0 mmol, 2.0 equiv.) were added and the flask was flushed with nitrogen, sealed, and stirred at 22 °C for 18 h. At this point reaction was judged complete, and the reaction mixture was diluted with ethyl acetate (150 mL) and washed with saturated sodium bicarbonate (2 x 100 mL), 10% citric acid in water (2 x 100 mL) and brine (1 x 100 mL). The combined organic phases were dried with anhydrous magnesium sulfate, filtered and the solvent removed *in vacuo*. The crude solid was recrystallized from ethanol affording N-[(2,4-dichlorophenyl)methyl]-4-nitrobenzamide (**1**) as pale yellow needles (3.72 g, 11.4 mmol, 95.6%).

**^1^H NMR** (400 MHz, DMSO-*d*_6_) δ 9.39 (t, *J* = 5.7 Hz, 1H), 8.37 – 8.31 (m, 2H), 8.17 – 8.11 (m, 2H), 7.64 (t, *J* = 1.2 Hz, 1H), 7.43 (d, *J* = 1.3 Hz, 2H), 4.55 (d, *J* = 5.7 Hz, 2H).

**^13^C NMR** (100 MHz, DMSO-*d*_6_) δ 165.32, 149.62, 140.00, 135.57, 133.51, 132.83, 130.83, 129.33, 129.11, 127.84, 124.06, 40.96.

**LC/MS** (ESI) exact mass for C_14_H_9_^35^Cl_2_N_2_O_3^-^_ [M-H]^-^ calc. 323.00, found 323.02.

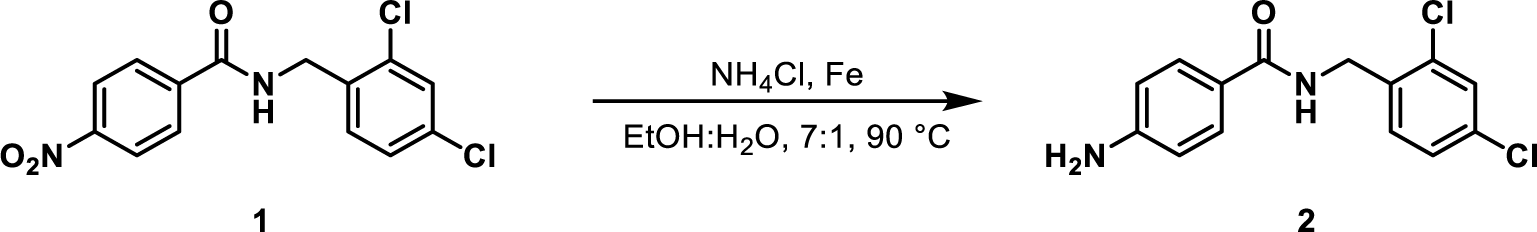

### Synthesis of 4-amino-N-(2,4-dichlorobenzyl)benzamide, 2

A round-bottom flask was charged with N-[(2,4-dichlorophenyl)methyl]-4-nitrobenzamide (**1**, 2.11 g, 6.49 mmol), iron powder (1.09 g, 19.5 mmol) and ammonium chloride (3.21 g, 64.9 mmol). Ethanol (100 mL) and water (15 mL) were added and the reaction was sealed and heated to 90 °C for 18 h. After cooling, the reaction mixture was filtered through diatomaceous earth, washing with ethyl acetate (25 mL). The combined filtrate was diluted with ethyl acetate (75 mL) and washed with dilute sodium hydroxide (2 x 100 mL) and brine (1 x 100 mL). The combined organics were dried with anhydrous magnesium sulfate, filtered and the solvent removed *in vacuo*. The crude solid was triturated with 25% ethyl acetate in hexanes and the remaining solid collected by vacuum filtration affording **2** as a pale tan solid (1.66 g, 5.61 mmol, 86.5% yield).

**^1^H NMR** (400 MHz, DMSO-*d*_6_) δ 8.60 (t, *J* = 5.9 Hz, 1H), 7.68 – 7.53 (m, 3H), 7.49 – 7.24 (m, 2H), 6.57 (d, *J* = 8.6 Hz, 2H), 5.67 (s, 2H), 4.45 (d, *J* = 5.8 Hz, 2H).

**_13_C NMR** (100 MHz, DMSO) δ 166.90, 152.36, 136.74, 133.13, 132.35, 130.31, 129.34, 128.89, 127.71, 120.96, 113.01.

**LCMS** (ESI) exact mass for C H ^35^Cl N O ^+^ [M+H]^+^ calcd: 295.04, found 294.91.

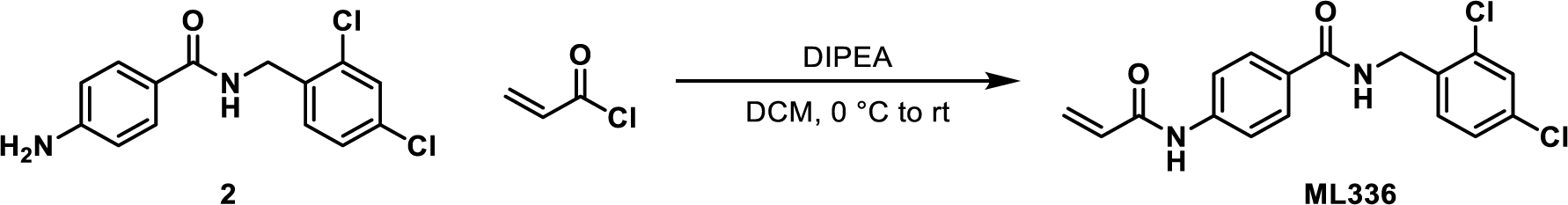

### Synthesis of ML336

A round-bottom flask was charged with 4-amino-N-(2,4-dichlorobenzyl)benzamide (**2**, 295 mg, 999 μmol, 1.00 equiv.), dichloromethane (5 mL) and diisopropylethylamine (348 μL, 2.00 mmol, 2.00 equiv.). The reaction was cooled to 0 °C and acryloyl chloride (109 mg, 1.20 mmol, 1.20 equiv) was added dropwise and the reaction allowed to warm to rt overnight. The crude reaction was concentrated *in vacuo*, then purified by column chromatography (0-100% EOAc in hexanes) affording ML336 as a white solid (201 mg, 576 μmol, 57.6%).

**^1^H NMR** (400 MHz, DMSO-*d*_6_) δ ppm 10.38 (s, 1H), 8.97 (t, J = 5.8 Hz, 1H), 7.91 (d, J = 8.8 Hz, 2H), 7.78 (d, J = 8.8 Hz, 2H), 7.62 (d, J = 2.1 Hz, 1H), 7.43 (dd, J = 8.4, 2.1 Hz, 1H), 7.37 (d, J = 8.4 Hz, 1H), 6.47 (m, 1H), 6.31 (dd, J = 16.9, 2.0 Hz, 1H), 5.81 (dd, J = 10.0, 2.0 Hz, 1H), 4.51 (d, J = 5.7 Hz, 2H).

**_13_C NMR** (100 MHz, DMSO) δ ppm 166.39, 163.90, 142.30, 136.23, 133.32, 132.58, 132.12, 130.53, 129.15, 129.01, 128.74, 127.98, 127.80, 119.08, 40.65, 40.44, 40.23, 40.02, 39.81, 39.60, 39.39.

**LCMS** (ESI) exact mass for C H ^35^Cl N O ^+^ [M+H]^+^ calcd: 349.05, found 348.99.

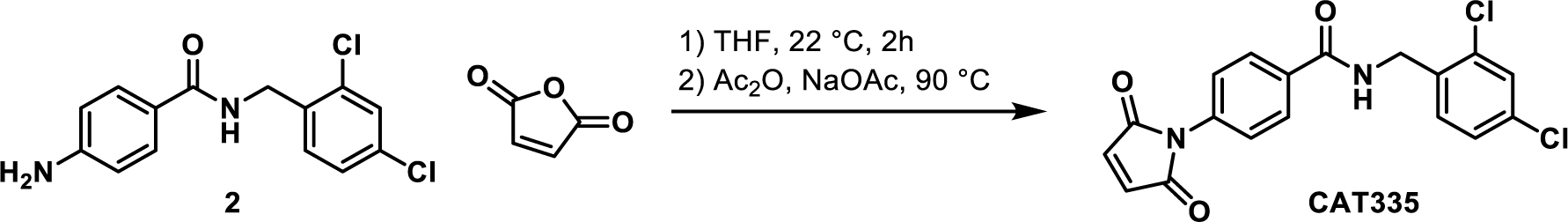

### Synthesis of CAT335

A vial was charged with 4-amino-N-(2,4-dichlorobenzyl)benzamide (**2**, 200 mg, 678 μmol, 1.00 equiv.), maleic anhydride (66.44 mg, 678 μmol, 1.00 equiv.) and anhydrous THF (1.5 mL). The reaction was flushed with nitrogen, sealed, and stirred at 22 °C. After 2 h, acetic anhydride (1.5 mL) and sodium acetate (167 mg, 2.03 mmol, 3.00 equiv.) were added and the reaction heated to 90 °C for 2 h. The reaction was cooled to rt, then diluted with water (10 mL). The resulting white precipitate was collected by vacuum filtration and washed with water (3 x 5 mL) and diethyl ether (3 x 5 mL), affording CAT335 as a white solid (200 mg, 533 μmol, 78.7%).

**^1^H NMR** (400 MHz, CDCl_3_) : δ ppm 7.90 (d, J = 7.7 Hz, 2H), 7.48-7.54 (m, 2H), 7.42-7.47 (m, 2H), 7.25-7.28 (m, 1H), 6.91 (s, 2H), 6.62 (br s, 1H), 4.72 ppm (d, J = 6.1 Hz, 2H).

**^13^C NMR** (100 MHz, CDCl_3_) δ ppm 168.99, 134.41, 134.37, 134.33 (br s, 1 C) 134.27, 134.08, 133.10, 131.36, 129.46, 127.87, 127.50, 125.61, 41.65.

**LCMS** (ESI) exact mass for C_14_H_9_^35^Cl_2_N_2_O_3^-^_ [M-H]^-^ calcd: 375.03, found 374.92.

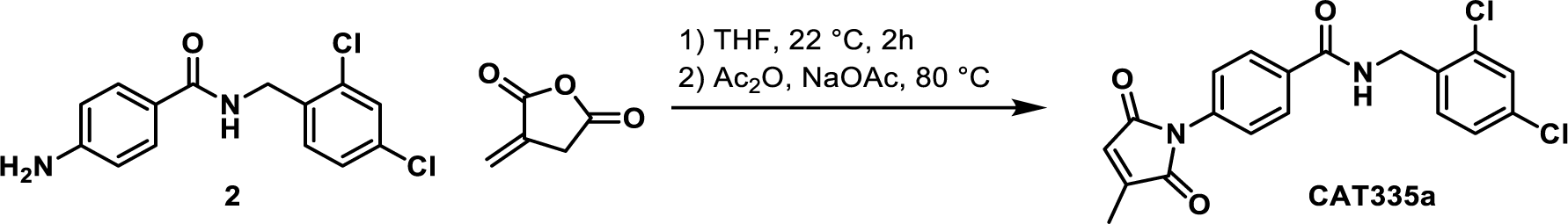

### Synthesis of CAT335a

A vial was charged with 4-amino-N-(2,4-dichlorobenzyl)benzamide (**2**, 30 mg, 102 μmol, 1.0 equiv.) and anhydrous THF (0.2 mL). Itaconic anhydride (13 mg, 116 μmol, 1.1 equiv.) was added and the reaction stirred for 30 min, after which a yellow precipitate formed. 0.2 mL acetic anhydride and sodium acetate (25 mg, . The reaction was flushed with nitrogen, sealed, and stirred at 22 °C. After 2 h, acetic anhydride (1.5 mL) and sodium acetate (25 mg, 3.0 mmol, 3.0 equiv.) were added and the reaction heated to 80 °C for 3 h. The reaction was cooled to rt, then quenched with sat. aq. NaHCO_3_ and extracted with EtOAc. The combined organics were dried with anhydrous sodium sulfate, filtered and the solvent removed *in vacuo*. The crude product was purified by column chromatography (0-100% EtOAc in hexanes) affording CAT335a as a white solid (18 mg, 46 μmol, 46%).

**^1^H NMR** (400 MHz, CDCl_3_) δ ppm 7.87 (d, J=8.64 Hz, 2 H) 7.48 (d, J=8.64 Hz, 2 H) 7.37-7.43 (m, 2 H) 7.23 (dd, J=8.28, 2.07 Hz, 1 H) 6.81 (br t, J=5.72 Hz, 1 H) 6.51 (q, J=1.70 Hz, 1 H) 4.68 (d, J=6.09 Hz, 2 H) 2.19 (d, J=1.70 Hz, 3 H).

**^13^C NMR** (100 MHz, CDCl_3_) δ ppm 170.18, 169.05, 166.58, 146.09, 134.77, 134.24, 134.16, 134.13, 132.76, 131.12, 129.39, 127.83, 127.68, 127.43, 125.36, 41.54, 11.20.

**LCMS** (ESI) exact mass for C_19_H_15_^35^Cl_2_N_2_O_3_^+^ [M+H]^+^ calcd: 389.05, found 389.14.

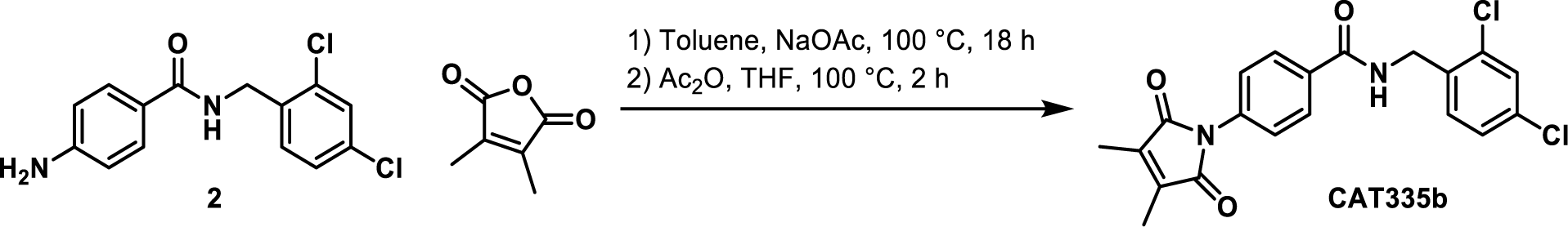

### Synthesis of CAT335b

A vial was charged with 4-amino-N-(2,4-dichlorobenzyl)benzamide (**2**, 59 mg, 200 μmol, 1.0 equiv.) and toluene (0.4 mL). 2,3-dimethyl maleic anhydride (25 mg, 200 μmol, 1.0 equiv.) and sodium acetate (49 mg, 600 μmol, 3.0 equiv.) were added and the reaction sealed, heated to 100 °C and stirred for 18 h. Acetic anhydride (0.4 mL) and THF (0.4 mL) were added and the reaction stirred for an additional 2 h. The reaction was cooled to rt, then quenched with sat. aq. NaHCO_3_ and extracted with EtOAc. The combined organics were dried with anhydrous sodium sulfate, filtered and the solvent removed *in vacuo*. The crude product was purified by column chromatography (0-100% EtOAc in hexanes) affording CAT335b as a white solid (25 mg, 62 μmol, 31%).

**^1^H NMR** (400 MHz, CDCl_3_) δ ppm 7.84-7.89 (m, 2 H) 7.49-7.53 (m, 2 H) 7.38 - 7.44 (m, 2 H) 7.24 (dd, J=8.28, 2.07 Hz, 1 H) 6.71 (br t, J=5.66 Hz, 1 H) 4.69 (d, J=5.97 Hz, 2 H) 2.08 (s, 6 H).

**^13^C NMR** (100 MHz, CDCl_3_) δ ppm 170.34, 166.10, 137.51, 135.73, 135.39, 133.53, 132.86, 132.78, 132.52,132.63, 130.50, 128.71 127.77, 127.22, 125.31, 40.64, 7.87.

**LCMS** (ESI) exact mass for C_20_H_17_^35^Cl_2_N_2_O_3_^+^ [M+H]^+^ calcd: 403.06, found 403.17.

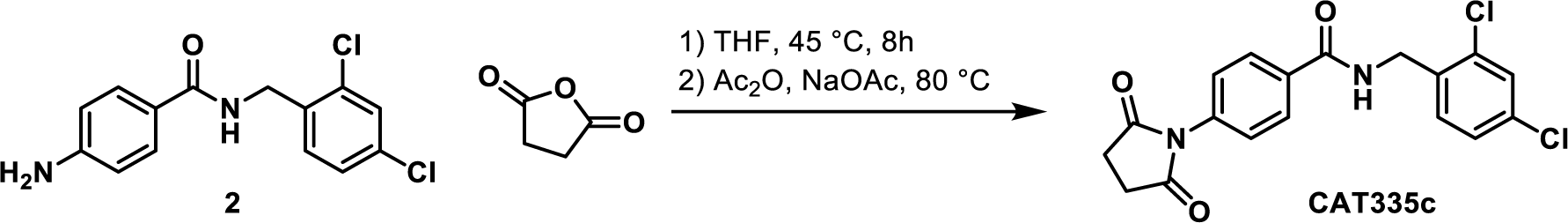

### Synthesis of CAT335c

A vial equipped with a magnetic stirbar was charged with 4-amino-N-(**2**, 4-dichlorobenzyl)benzamide (**2**,148 mg, 500 µmol, 1 equiv.), succinic anhydride (50 mg, 500 µmol, 1 equiv.) and anhydrous THF (2 mL) and heated to 45 °C for 8 h. Acetic anhydride (2 mL) and sodium acetate (123 mg, 1.50 mmol, 3 equiv.) were then added and the reaction heated to 80 °C for 16 h. The reaction was cooled to rt and diluted with water (5 mL). The resulting white solid was collected by vacuum filtration, then recrystallized from 3:1 Ethyl acetate:hexanes to afford CAT335c as a white solid (82.0 mg, 217 µmol, 43.5%).

**^1^H NMR** (400 MHz, DMSO-*d*_6_) δ = 9.16 (br t, *J* = 5.7 Hz, 1H), 8.02 (d, *J* = 8.0 Hz, 2H), 7.64 (s, 1H), 7.47 - 7.36 (m, 4H), 4.54 (d, *J* = 5.6 Hz, 2H), 2.81 (s, 4H).

**^13^C NMR** (100 MHz, DMSO-*d*_6_) δ = 177.2, 166.3, 136.0, 135.8, 134.0, 133.4, 132.7, 130.6, 129.1, 128.4, 127.8, 127.4, 40.8, 29.0.

**LCMS** (ESI) exact mass for C_18_H_15_^35^Cl_2_N_2_O_3_^+^ [M+H]^+^ calcd: 377.05, found 376.91.

