## Supplementary Figs. S1-S8, Scheme 1, Tables S1-S4, and references for "Development of covalent chemogenetic K_2P_ channel activators"

15 October 2023

**Supplementary Material for**
**Development of covalent chemogenetic $K_{2P}$ channel activators**

Parker E. Deal [1,2], Haerim Lee[1], Abhisek Mondal[1], Marco Lolicato[1#], Philipe Ribeiro Furtado de Mendonca[3], Holly Black[3], Xochina El-Hilali[2], Clifford Bryant[2], Ehud Y. Isacoff[3-6], Adam R. Renslo[2]*, and Daniel L. Minor Jr.[1,6-9]*

[1]Cardiovascular Research Institute
[2]Department of Pharmaceutical Chemistry
[7]Departments of Biochemistry and Biophysics, and Cellular and Molecular Pharmacology
[8]California Institute for Quantitative Biomedical Research
[9]Kavli Institute for Fundamental Neuroscience
University of California, San Francisco, California 93858-2330 USA

[3]Department of Molecular and Cell Biology,
[4]Helen Wills Neuroscience Institute
[5]Weill Neurohub
University of California, Berkeley, Berkeley, California 94720, United States.

[6]Molecular Biophysics and Integrated Bio-imaging Division
Lawrence Berkeley National Laboratory, Berkeley, CA 94720 USA

### Current address: Department of Molecular Medicine, University of Pavia, Pavia, Italy 27100

#### Figure S1 — Deal *et al.*

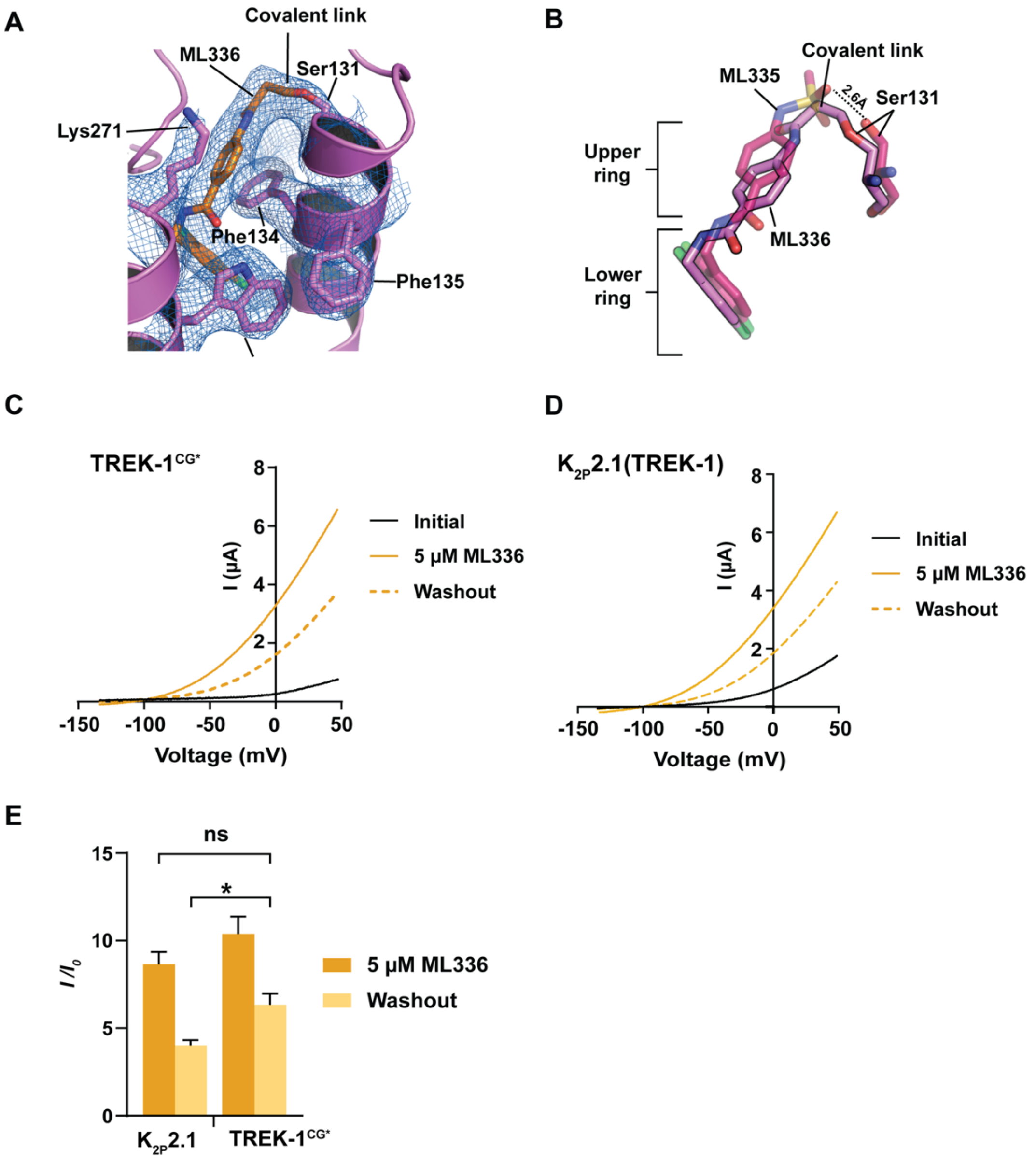

**Figure S1 ML336 structure and functional analysis A,** Exemplar 2.9Å resolution 2*Fo-Fc* electron density (1σ) showing the $K_{2P}$2.1(TREK 1):ML336 covalent complex. Select residues are indicated. $K_{2P}$2.1 sidechains are magenta. ML336 is orange. Covalent link is indicated. P1 and M4 helices are labeled. **B,** Activators and Ser131 residues from the $K_{2P}$2.1(TREK-1):ML336 (magenta) and $K_{2P}$2.1(TREK-1):ML335 (PDB:6CQ8)(pink) {Lolicato, 2017 #1406} complexes. **C,** and **D,** Exemplar two-electrode voltage clamp (TEVC) traces from *Xenopus* oocytes expressing **C,** TREK-$1^{CG*}$ and **D,** $K_{2P}$2.1(TREK-1) showing baseline, 2 minute application of 5 μM ML336, and after 2 minutes of washout. **E,** Activation ($I/I_0$) of $K_{2P}$2.1(TREK-1) and TREK-$1^{CG*}$ currents at 0 mV following application of 5 μM ML336 for 2 min and after washout (2 minutes buffer) (n=13-16). Significance measured by two-sided, unpaired, unequal variances t-test where: ns = $p>0.05$, * = $p<0.05$. Error bars show S.E.M

**Figure S2** **Deal *et al.***

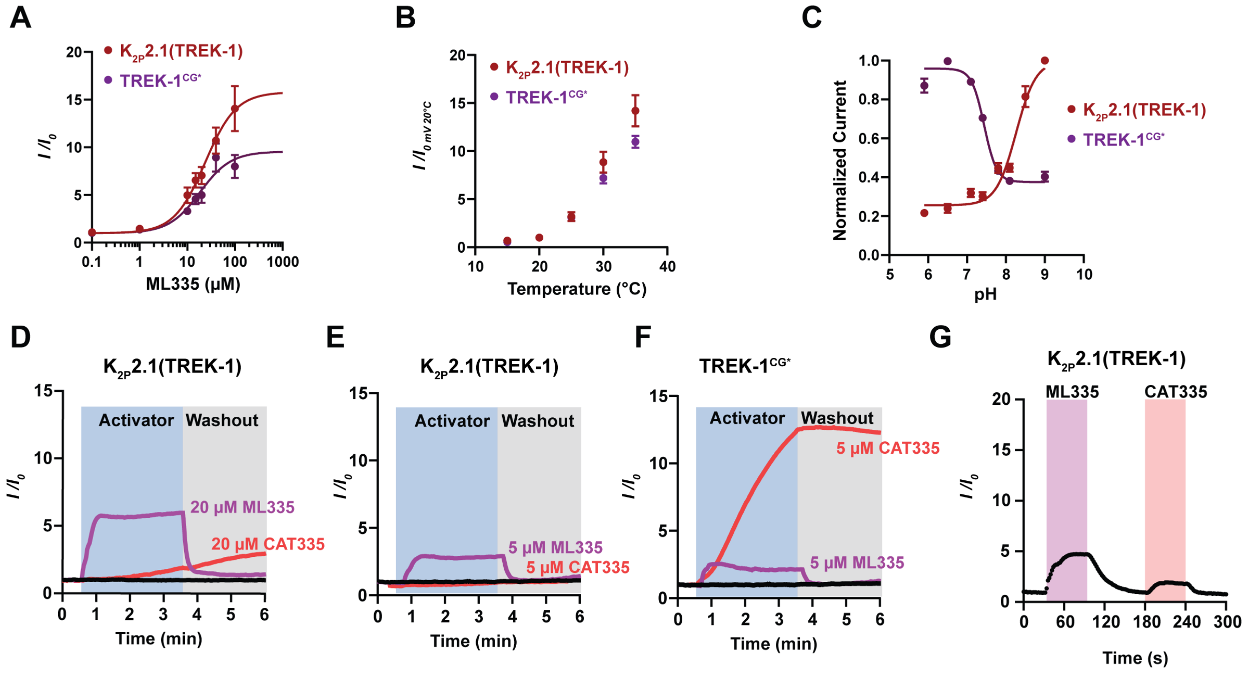


**Figure S2 Comparison of TREK-1$^{CG*}$ and K2P2.1(TREK-1) functional properties A-C,** Comparison of **A,** ML335 dose-response, **B,** Temperature response, and **C,** Extracellular pH responses for K2P2.1(TREK-1) (red) and TREK-1$^{CG*}$ (purple) measured by TEVC in *Xenopus* oocytes. For '**A**' $EC_{50}$= 18 ± 4 and 24 ±4 µM; $E_{max}$ = 9.6 ± 0.8, 15.8 ± 1.3 for K2P2.1(TREK-1) and TREK-1$^{CG*}$, respectively. Currents were measured by TEVC at 0 mV and normalized to initial currents before ML335 application (n=16-19). For '**B**', Currents were normalized to the current at 20 °C and 0 mV for each channel (n=10). For '**C**' Currents were measured by TEVC at 0 mV K2P2.1(TREK-1) is inhibited at acidic pH (pK=8.3 ± 0.1, Hillslope=1.7). TREK-1$^{CG*}$ is activated at acidic pH (pK=7.4 ± 0.1, Hillslope=-2.7 ± 0.4). Currents were normalized to the highest mean current for each channel (n=9-21). **D-F,** Representative TEVC time courses of **D,** and **E,** K2P2.1(TREK-1), and **F,** TREK-1$^{CG*}$ activation (*$I/I_0$*) with ML335 (magenta) CAT335 (red) at 5 µM or 20 µM concentrations. **G,** Exemplar time course of K2P2.1(TREK-1) activation in HEK293 cells by 20 µM ML335 (magenta) and 20 µM CAT335 (red). A' was fit to the Hill equation Y= 1+(X^HillSlope)*(Max-1)/(X^Hillslope+ $EC_{50}$^HillSlope). 'C' was fit to the Hill equation Y=Bottom + (1-Bottom)/(1+10^((Log$EC_{50}$-X)*HillSlope))). Error bars are ± S.E.M

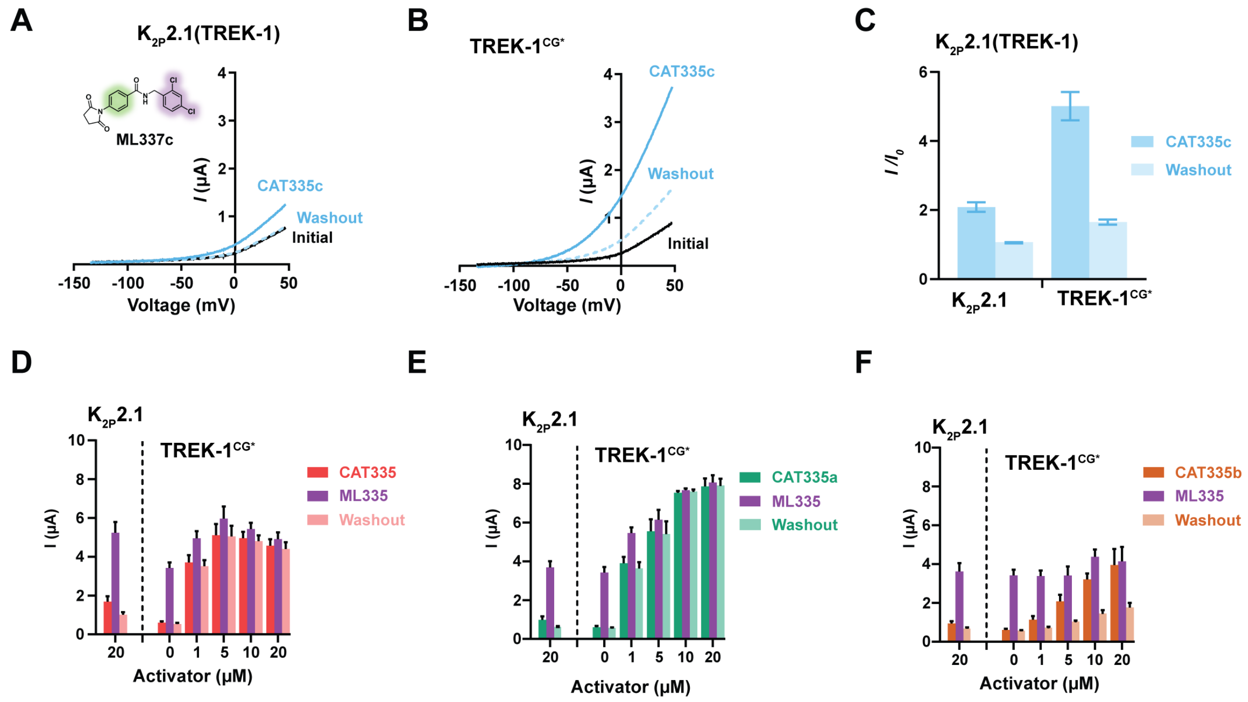


**Figure S3 CAT335 derivative functional properties. A-B**, Exemplar TEVC recordings of *Xenopus* oocytes expressing **A,** $K_{2P}2.1$(TREK-1) or **B,** TREK-$1^{CG*}$ in response to 20 µM CAT335c. **C,** Activation ($I/I_0$) of $K_{2P}2.1$(TREK-1) and TREK-$1^{CG*}$ currents at 0 mV following application of 20 µM CAT335c for 2 min and after 2 min washout (n=9). **D-F,** TEVC current magnitude at 0 mV recorded from oocytes expressing $K_{2P}2.1$(TREK-1) or TREK-$1^{CG*}$ following 1 hour incubation with **D,** CAT335, **E,** CAT335a, or **F,** CAT335b. After recording the initial currents following covalent activator incubation, oocytes were treated with 20 µM ML335. Following ML335 activation, the oocytes were perfused with buffer for 3 minutes to measure washout. Dashed lines separate $K_{2P}2.1$(TREK-1) and TREK-$1^{CG*}$ responses. Error bars are ± S.E.M..

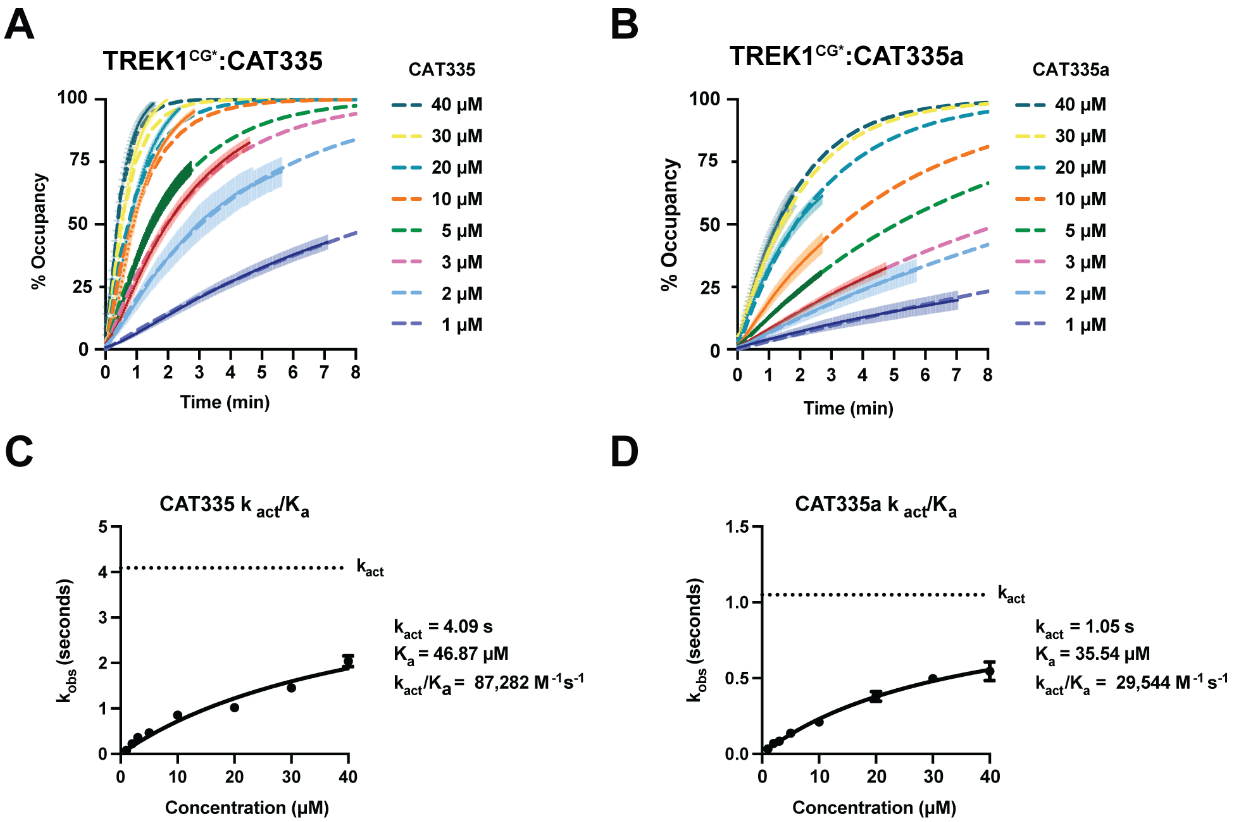


**Figure S4 Kinetic analysis of CAT335 and CAT335a activation of TREK-1$^{CG*}$. A-B,** TEVC recordings from *Xenopus* oocytes expressing TREK-1$^{CG*}$ treated with 1-40 µM of **A,** CAT335 or **B,** CAT335a. % Occupancy was defined by the maximal activation observed following the addition of 40 µM CAT335 for 4 minutes at the end of each time course. Each time course was fitted using the Y=100(1-exp(-$k_{obs}$*X))( n=3). Confidence intervals represent S.E.M.. **C-D,** The $k_{obs}$ values were plotted vs. concentration and fitted to the equation Y = ($K_{act}$*X)/($K_a$+X).

### Figure S5

Deal *et al.*

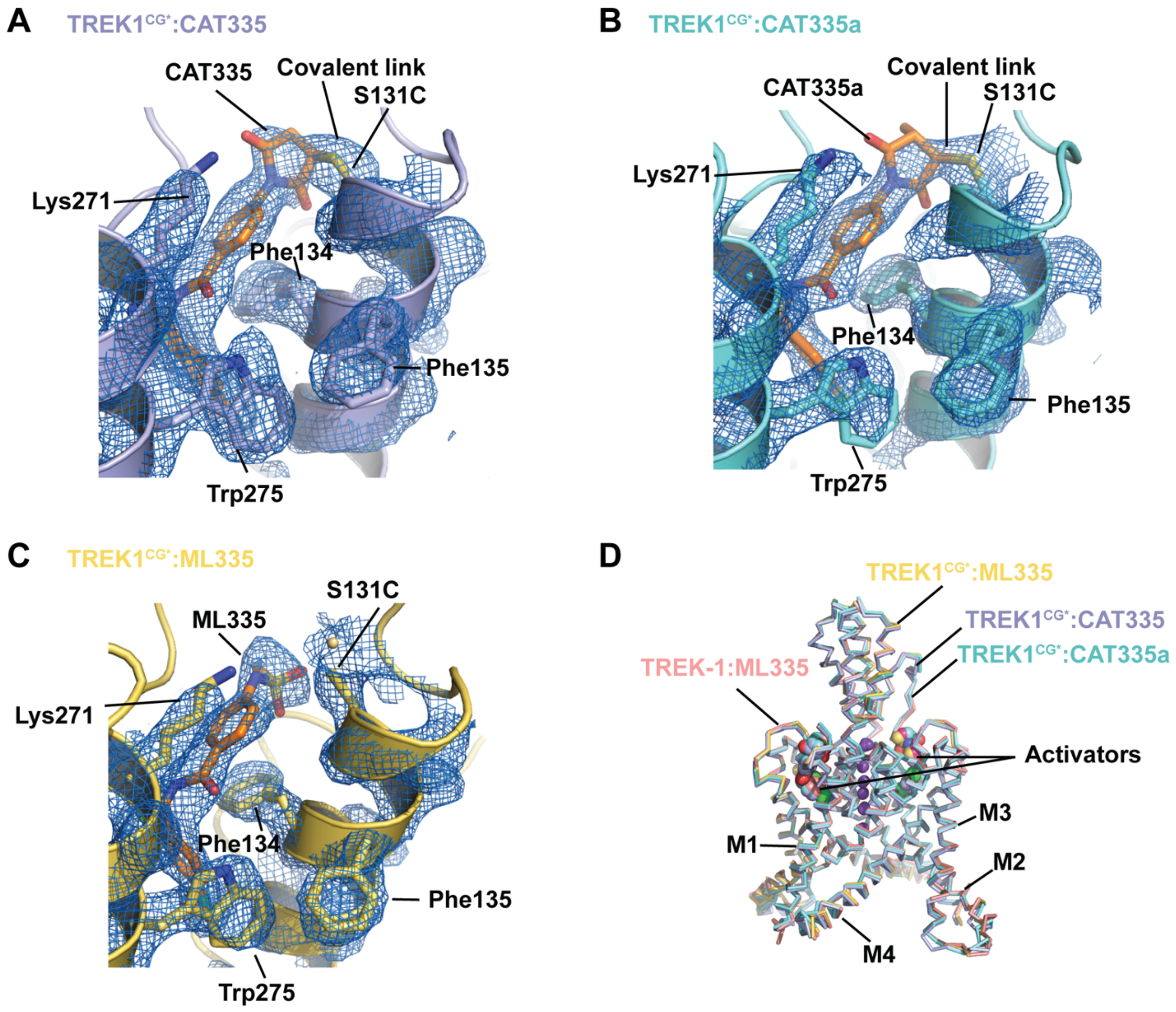


**Figure S5 Structural analysis of chemo-genetic pairs.** Exemplar 2*Fo*-*Fc* electron density (1σ) for **A,** TREK-1$^{CG*}$:CAT335 (light blue), **B,** TREK-1$^{CG*}$:CAT335a (aquamarine), and **C,** TREK-1$^{CG*}$:ML335 (yellow) complexes. Select residues are indicated. CAT335, CAT335a, and ML335 are orange. Covalent links are indicated. **D,** Superposition of TREK-1$^{CG*}$:CAT335 (light blue), TREK-1$^{CG*}$:CAT335a (aquamarine), TREK-1$^{CG*}$:ML335 (yellow), and $K_{2P}$2.1(TREK-1):ML335 (PDB:6CQ8){Lolicato, 2017 #1406} (pink) complexes. Activators are shown in space filling. Potassium ions are purple spheres.

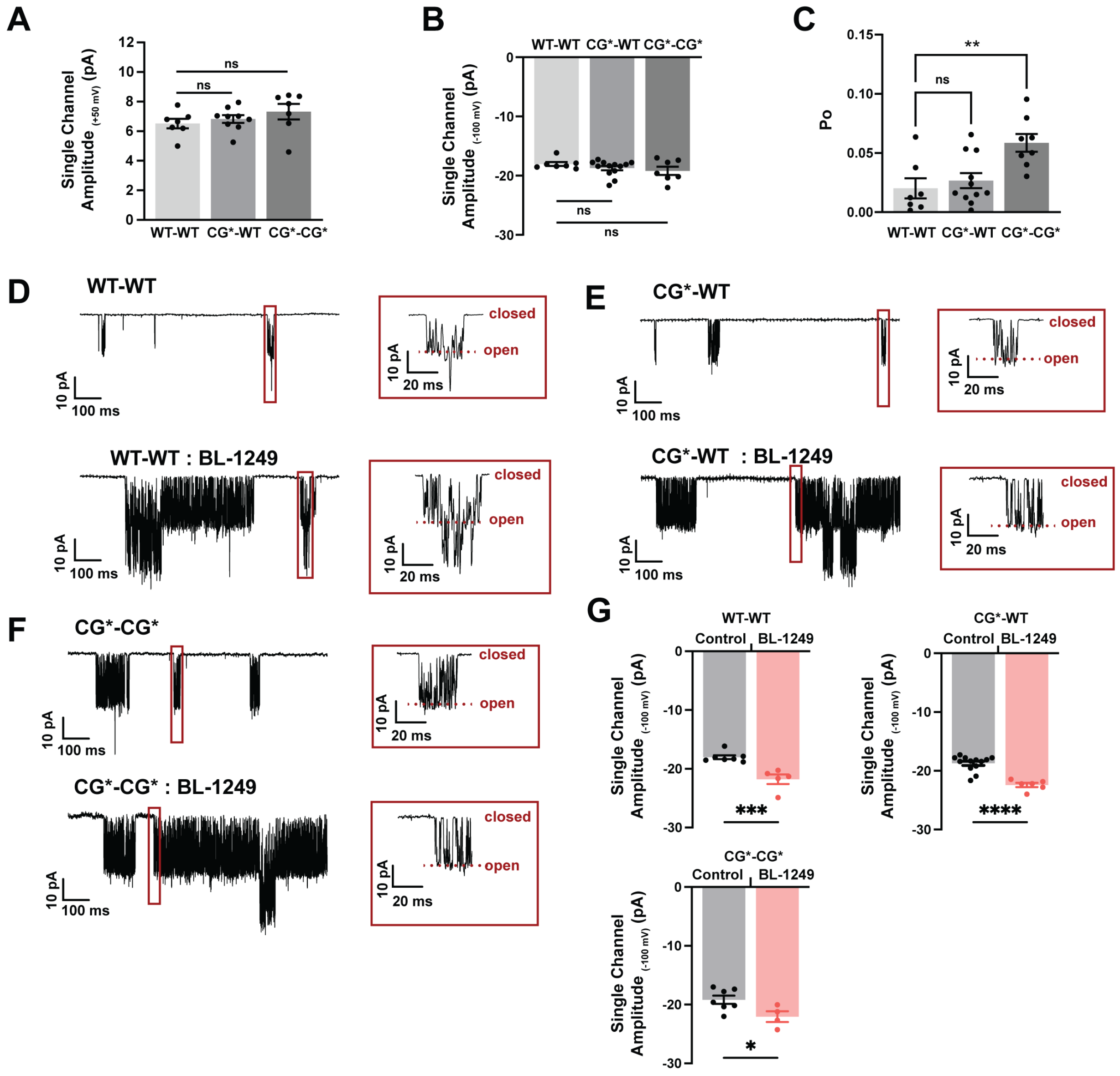
A
Single Channel Amplitude (+50 mV) (pA)
ns
ns
WT-WT CG*-WT CG*-CG*
B
WT-WT CG*-WT CG*-CG*
Single Channel Amplitude (-100 mV) (pA)
ns
ns
C
Po
**
ns
WT-WT CG*-WT CG*-CG*
D
WT-WT
closed
open
10 pA
100 ms
20 ms
WT-WT : BL-1249
E
CG*-WT
CG*-WT : BL-1249
F
CG*-CG*
CG*-CG* : BL-1249
G
WT-WT
Control BL-1249
***
CG*-WT
****
CG*-CG*
*

**Figure S6 $K_{2P}$2.1(TREK-1) tandem construct single channel analysis**. **A-B,** Basal single channel amplitude at **A**, +50 mV and **B**, -100 mV for the indicated tandem constructs. **C,** Basal $P_o$ for the indicated tandem constructs. **D-F,** Exemplar cell attached single channel recordings at -100 mV for **D,** WT-WT, **E,** CG*-WT, and **F,** CG*-CG* (top) and after treatment with 20 µM BL-1249 (bottom). Insets in D-F show boxed regions. **G,** Single channel amplitude at -100 mV for the indicated channels under basal (grey) and 20 µM BL-1249 (red) conditions. Significance for (A-C) was measured using Brown-Forsythe and Welch one-way ANOVA with Dunnett's T3 multiple comparisons test (n.s. p>0.12; **p<0.0021). Significance for panel G was measured using unpaired t-tests (n.s. p>0.12; * p<.033; ** p<.0021; *** p<.0002; **** p<.0001),

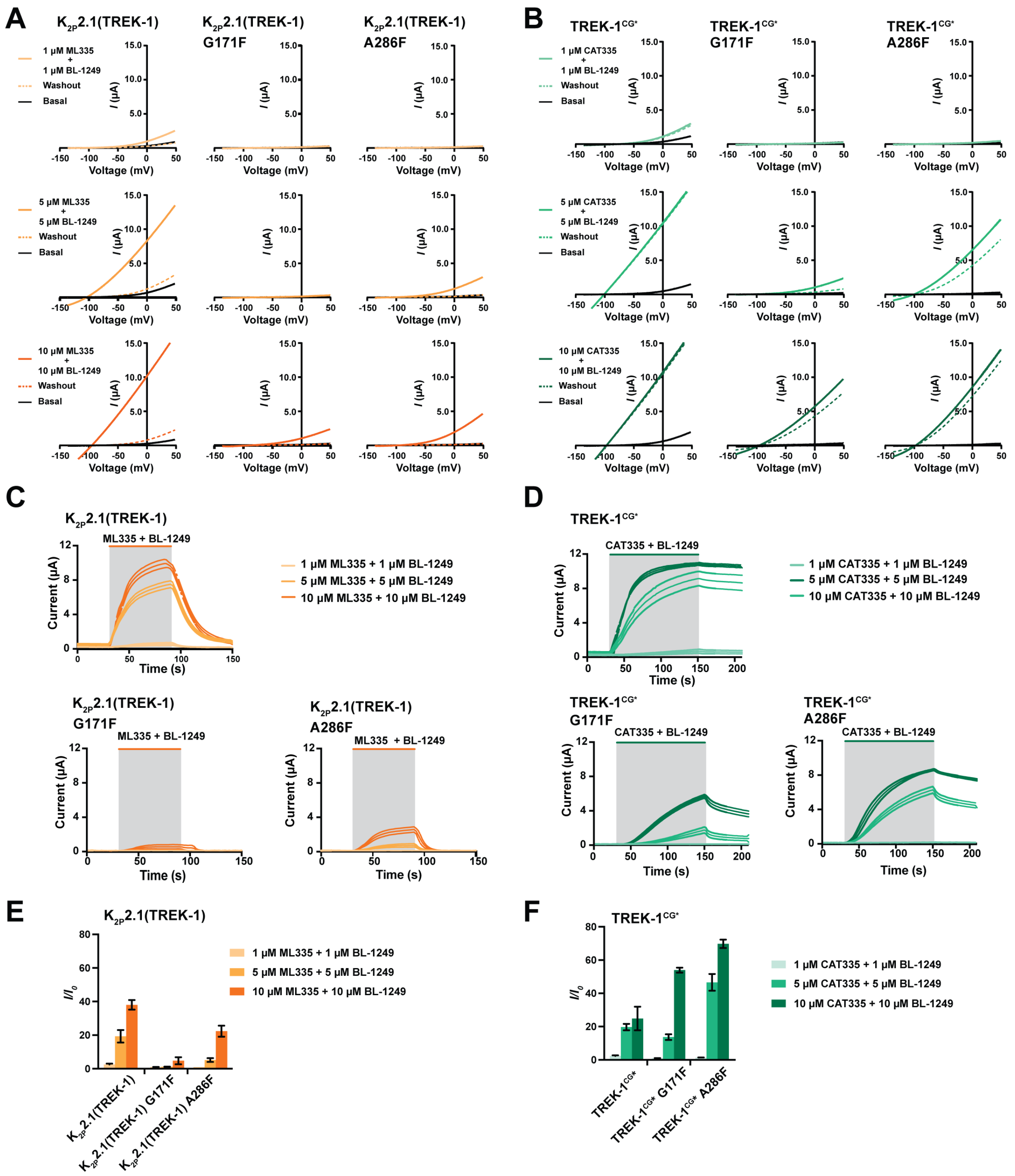
A
K2P2.1(TREK-1)
K2P2.1(TREK-1) G171F
K2P2.1(TREK-1) A286F
1 μM ML335 + 1 μM BL-1249
5 μM ML335 + 5 μM BL-1249
10 μM ML335 + 10 μM BL-1249
Washout
Basal
I (μA)
Voltage (mV)
B
TREK-1CG*
TREK-1CG* G171F
TREK-1CG* A286F
1 μM CAT335 + 1 μM BL-1249
5 μM CAT335 + 5 μM BL-1249
10 μM CAT335 + 10 μM BL-1249
C
ML335 + BL-1249
Current (μA)
Time (s)
D
CAT335 + BL-1249
E
I/I0
F

**Figure S7 $K_{2P}$2.1(TREK-1) and TREK-1$^{CG*}$ responses to modulator pocket ligand (ML335 or CAT335) and fenestration site ligand BL-1249 co-application. A,** Exemplar TEVC traces from *Xenopus* oocytes expressing $K_{2P}$2.1(TREK-1) (left), $K_{2P}$2.1(TREK-1) G171F (center), and $K_{2P}$2.1(TREK-1) A286F (right) showing responses to simultaneous application of ML335 and BL-1249 at the indicated concentrations and after washout (dashed lines). **B,** Exemplar TEVC traces from *Xenopus* oocytes expressing TREK-1$^{CG*}$ (left), TREK-1$^{CG*}$ G171F (center), and TREK-1$^{CG*}$ A286F (right) showing responses to the simultaneous application of CAT335 and BL-1249 at the indicated concentrations and after washout (dashed lines). **C,** Activation timecourses at 0 mV for $K_{2P}$2.1(TREK-1), $K_{2P}$2.1(TREK-1) G171F, and $K_{2P}$2.1(TREK-1) A286F to simultaneous application of the indicated concentrations of ML335 and BL-1249 (grey shading). **D,** Activation timecourses at 0 mV for TREK-1$^{CG*}$, TREK-1$^{CG*}$ G171F, and TREK-1$^{CG*}$ A286F to simultaneous application of the indicated concentrations of ML335 and BL-1249 (grey shading). **E,** Activation ($I/I_0$) for $K_{2P}$2.1(TREK-1), $K_{2P}$2.1(TREK-1) G171F, and $K_{2P}$2.1(TREK-1) A286F for the indicated activator concentrations. **F,** Activation ($I/I_0$) of TREK-1$^{CG*}$, TREK-1$^{CG*}$ G171F , and TREK-1$^{CG*}$ A286F for the indicated activator concentrations. Error bars are ± S.E.M..

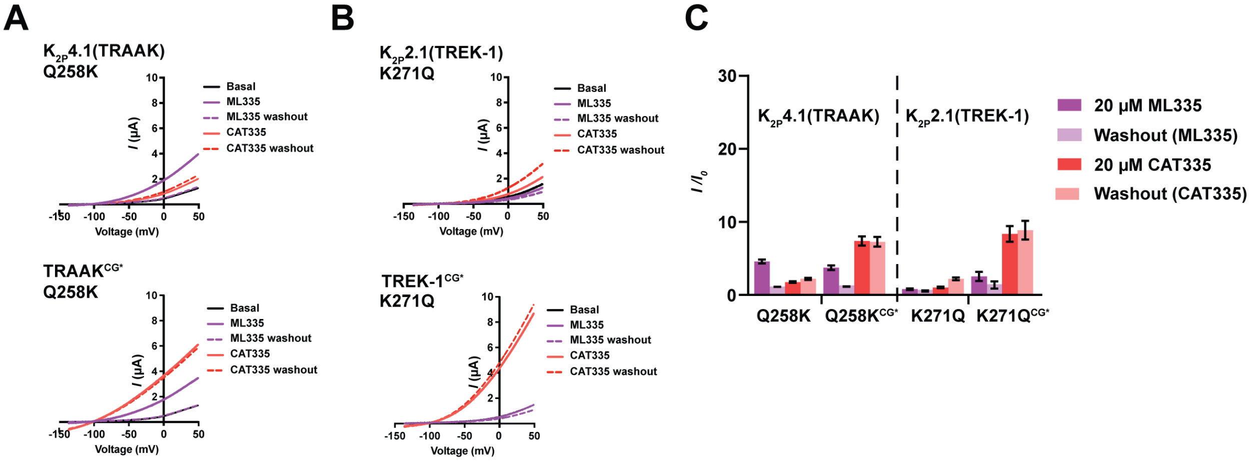


**Figure S8 $K_{2P}$ Modulator Pocket Cation-π interaction affects CAT335 responses. A-B,** Exemplar TEVC recordings of **A,** $K_{2P}$2.1(TREK-1) K271Q (top) and TREK-1$^{CG*}$ K271Q (bottom), and **B,** $K_{2P}$4.1(TRAAK) Q258K (top) and $K_{2P}$4.1(TRAAK)$^{CG*}$ Q258K (bottom) to 2 minute application and washout of 20 µM ML335 and 20 µM CAT335 in series. **C,** Activation ($I/I_0$) at 0 mV of *Xenopus* oocytes expressing the indicated $K_{2P}$ channels following application of 20 µM ML335 (magenta) or 20 µM CAT335 (red) (n=3-10). Error bars are ± S.E.M..

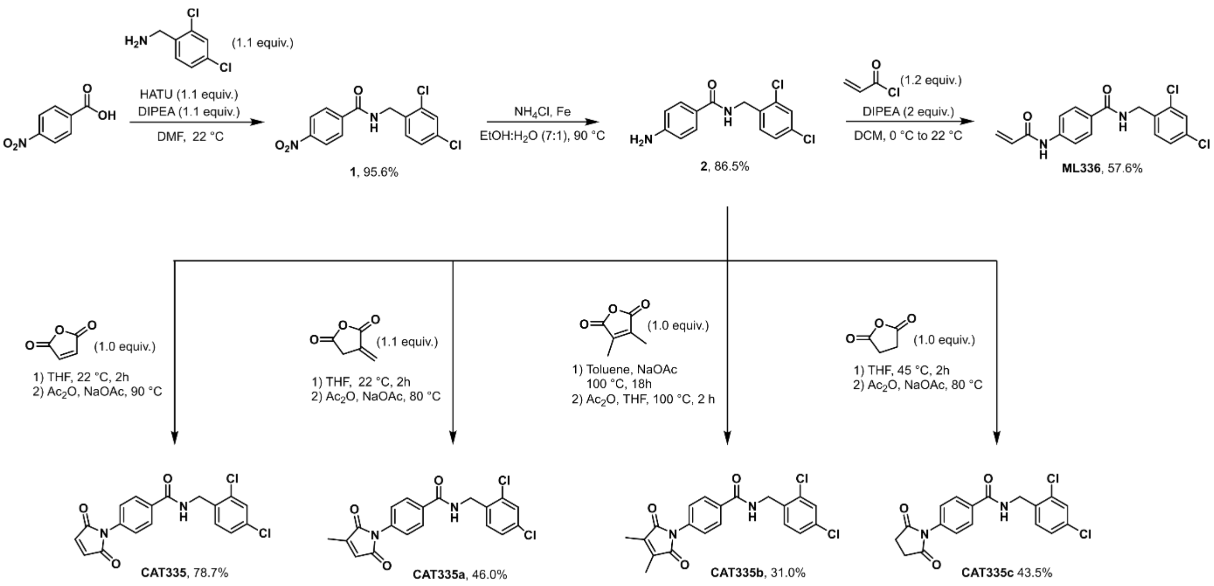


**Scheme 1: Synthesis of ML336 and CAT335 derivatives**

**Table S1 Crystallographic data collection and refinement statistics**

| | **$K_{2P}$2.1 (TREK-1):ML336 (PDB:8UF6)** | **TREK-1$^{CG*}$:CAT335 (PDB: 8UE9)** | **TREK-1$^{CG*}$:CAT335a (PDB:8UEC)** | **TREK-1$^{CG*}$:ML335 (PDB:8UE2)** |
|---|---|---|---|---|
| **Data Collection** | | | | |
| Space group | P $2_12_12_1$ | P $2_12_12_1$ | P $2_12_12_1$ | P $2_12_12_1$ |
| Cell dimensions a/b/c (Å) | 66.953/118.597/130.873 | 67.07/119.78/129.71 | 67.213/120.568128.613 | 67.213/120.568128.613 |
| α/β/γ (°) | 90/90/90 | 90/90/90 | 90/90/90 | 90/90/90 |
| Resolution (Å) | 14.82 - 2.901 (3.004 - 2.901) | 46.6 - 3.0 (3.107 - 3.0) | 29.35 - 3.0 (3.107 - 3.0) | 24.53 - 3.0 (3.107 - 3.0) |
| $R_{merge}$ (%) | 0.2041 (>1) | 0.290(>1) | 0.222(>1) | 0.232(>1) |
| I / σ I | 8.73 (0.42) | 6.8(0.3) | 6.5(0.3) | 10.9 (0.7) |
| CC(1/2) | 0.997 (0.209) | 0.993 (0.207) | 0.998(0.145) | 0.998 (0.261) |
| Completeness (%) | 97.97 (94.39) | 99.85 (99.81) | 100 (99.3) | 100 (99.4) |
| Redundancy | 7.2 (7.3) | 19.0 (15.5) | 12.9 (8.0) | 17.1 (9.3) |
| Total reflections | 168711 (16472) | 43150 (4232) | 43155 (4224) | 43140 (4218) |
| Unique reflections | 23483 (2260) | 21576 (2112) | 21579 (1555) | 21571 (1552) |
| Wilson B-factor | 91.74 | 118.14 | 107.74 | 107.74 |
| Wavelength (Å) | 0.9779 | 0.9779 | 0.9779 | 0.9779 |
| **Refinement** | | | | |
| $R_{work}$ / $R_{free}$ (%) | 26.7/ 31.3 | 26.0/0.30.6 | 27.3/29.3 | 26.6/29.2 |
| No. of chains in AU | 2 | 2 | 2 | 2 |
| No. of protein atoms | 4273 | 4340 | 4280 | 4357 |
| No. of ligand atoms | 210 | 302 | 257 | 224 |
| No. of water atoms | 0 | 1 | 0 | 0 |
| RMSD bond lengths (Å) | 0.006 | 0.002 | 0.004 | 0.004 |
| RMSD angles (°) | 1.15 | 0.54 | 0.66 | 0.69 |
| Ramachandran favored/allowed/outliers (%) | 92.46/7.17/0.37 | 95.64/3.45/0.91 | 92.14/6.67/1.1 | 94.02/4.89/1.09 |

#### Table S2 Activator responses

| | | ML335* | | ML336 | | CAT335a | | CAT335b | | CAT335c | |
|---|---|---|---|---|---|---|---|---|---|---|---|
| | | *I/I₀* | n | *I/I₀* | n | *I/I₀* | n | *I/I₀* | n | *I/I₀* | n |
| $K_{2P}$2.1 (TREK-1) | WT | 6.5 ± 0.3 | 10 | 10.0 ± 1.0 | 11 | 1.7 ± 0.1 | 8 | 1.3 ± 0.1 | 6 | 2.1 ± 0.1 | 9 |
| | | - | - | 8.7 ± 0.7** | 16 | - | - | - | - | - | - |
| | S131A | 6.8 ± 0.7 | 6 | 16.1 ± 2.5 | 6 | - | - | - | - | - | - |
| | S131C (CG*) | 3.6 ± 0.4 | 10 | 10.4 ± 1.0** | 13 | 7.4 ± 0.7 | 8 | 4.9 ± 0.6 | 9 | 5.0 ± 0.4 | 9 |

Values are for 20µM except where noted.

*ML335 data are identical to those in Table 1.
**Activation by 5 µM ML336

**Table S3 Responses to simultaneous or sequential MP and FS activator application.**

| | | Simultaneous | | | | Sequential | | | | | | | |
|---|---|---|---|---|---|---|---|---|---|---|---|---|---|
| | | 10 μM ML335 + 10 μM BL1249 | | 10 μM CAT335 + 10 μM BL1249 | | 1) 20 μM CAT335 | | 2) 20 μM BL-1249 | | 1) 20 μM BL-1249 | | 2) *20 μM ML335*† Or 20 μM CAT335 | |
| | | *I/I₀* | n | *I/I₀* | n | *I/I₀* | n | *I/I₀* | n | *I/I₀* | n | *I/I₀* | n |
| $K_{2P}2.1$ (TREK-1) | WT | 38 ± 3 | 3 | - | - | - | - | - | - | 15.4 ± 1.7 | 4 | 18.9 ± 1.2† | 4† |
| | G171F | 4.7 ± 2 | 3 | - | - | 1.02 ± 0.02 | 5 | 3.8 ± 0.5 | 5 | 4 ± 3 | 4 | 3.5 ± 1.2† | 4† |
| | A286F | 22 ± 4 | 3 | - | - | 1.03 ± 0.03 | 7 | 19.4 ± 1.3 | 7 | 9.4 ± 0.9 | 6 | 10.1 ± 1.0† | 6† |
| | CG* | - | - | 25 ± 7 | 3 | - | - | - | - | 12.0 ± 1.6 | 3 | 15 ± 2 | 3 |
| | CG*/G171F | - | - | 54 ± 1 | 3 | 7.8 ± 1.3 | 9 | 62 ± 3 | 9 | 10 ± 2 | 3 | 21 ± 2 | 3 |
| | CG*/A286F | - | - | 70 ± 3 | 3 | 21 ± 3 | 8 | 53 ± 5 | 8 | 10.3 ± 1.2 | 3 | 26 ± 3 | 3 |

#### Table S4 Hippocampal neuron responses to CAT335

| | WT (n = 19) | | CG* A286F (n=11) | |
|---|---|---|---|---|
| | **Baseline** | **20 µM CAT335** | **Baseline** | **20 µM CAT335** |
| **Membrane potential (mV)** | -65.5 ± 0.7 | -68.5 ± 0.9 | -68.7 ± 1.8 mV | -80.7 ± 2.2 |
| **Δ Membrane potential (%)** | n.a. | 105% | n.a. | 118% |
| **Input resistance (MΩ)** | 294 ± 31 | 290 ± 31 | 253 ± 54 | 153 ± 56 |
| **Δ Input resistance (%)** | n.a. | 99 | n.a. | 57 |

n.a. – not applicable
